# Model-Agnostic Decomposition of the Difference in Mean Response Between Two Groups via Game-Theoretic Attribution with Minimum-Switching-Distortion Coupling

**DOI:** 10.64898/2026.11.02.763621

**Authors:** Hyunwook Koh

## Abstract

The difference in mean response between two groups can arise from differences in covariate–response relationships, covariate distributions, or both. We propose a model-agnostic decomposition (MAD) of this difference into structure and composition effects. We formulate a cooperative switching game with a model-switch player and individual covariate-switch players, organized into model and covariate unions. MAD employs the Owen value, whose quotient game yields the aggregate structure and composition effects and whose internal game allocates the aggregate composition effect across covariates, and thereby inherits its fair allocation properties. In the internal game, partial covariate switches create hybrid covariate vectors whose joint distributions are not determined by the group-specific covariate distributions alone. To resolve this indeterminacy, we propose a minimum-switching-distortion (MSD) coupling, formulated as an optimal-transport problem. The MSD coupling satisfies admissibility conditions of identity at coincidence, transposition equivariance, and coordinate equivariance, and its sparse support reduces allocation cost. Because MAD imposes no particular functional form, it accommodates machine-learning models that adapt to complex nonlinear and interactive relationships, thereby yielding more accurate decompositions. Simulation studies demonstrate the effectiveness of MAD and the MSD coupling in terms of decomposition accuracy, coupling admissibility, and computational cost. As an illustration, we decompose the difference in mean body mass index between UK and US males aged 30–70 years using microbiome data, at the aggregate level and across microbial genera.

## 1 Introduction

The difference in mean response between two groups is of interest in many areas of economics, social science, epidemiology, and biomedical research. A central question is how much of the mean-response difference is associated with differences in covariate composition and how much with differences in the covariate–response relationship. The Blinder–Oaxaca decomposition (Blinder, 1973; Oaxaca, 1973) addresses this question under linear regression models by decomposing the mean-response difference into a composition effect due to differences in covariate means and a structure effect due to differences in regression coefficients. The linearity and additivity of the model also permit straightforward covariate-specific decompositions of the composition and structure effects.

A substantial literature has extended the Blinder–Oaxaca decomposition beyond linear regression models. Yun (2004) and Sinning et al. (2008) developed decomposition methods for nonlinear models, while Rothe (2015) further decomposed the composition effect under general nonlinear relationships into contributions from marginal covariate distributions, their dependence, and their interactions. More recently, Mao et al. (2025) developed a double-machine-learning approach for estimation and inference on Oaxaca–Blinder counterfactual means and aggregate effects. We likewise accommodate nonlinear and interactive predictive models, including black-box machine-learning models, without requiring a coefficient-based representation. In addition, we leverage game-theoretic attribution to ensure desirable allocation properties.

Game-theoretic attribution provides a principled way to address order dependence in decomposition problems. The Shapley value (Shapley, 1953) offers a general principle for allocating the total value of a cooperative game across its players according to their average marginal contributions and is uniquely characterized by the axioms of efficiency, symmetry, additivity, and the null-player property. Owen (1977) extended the Shapley value to games with *a priori* unions, in which players are partitioned into unions, allowing contributions to be allocated first across unions and then across players within each union. Shorrocks (2013) subsequently developed a unified game-theoretic framework for decomposition analysis, including hierarchical decompositions based on grouped contributory factors, while Audoly et al. (2025) more recently extended this line of work to nonlinear economic models. By averaging marginal contributions across possible orderings, these approaches yield order-invariant, exactly additive allocations under nonlinear relationships.

Other Shapley-based methods have also addressed related problems involving distribution shift and predictive performance. Zhang et al. (2023) used Shapley values to decompose predictive-performance differences across environments into contributions from shifts in the underlying data-generating mechanisms, while Singh et al. (2024) hierarchically decomposed predictive-performance differences across domains into distribution-shift components and their Shapley-based covariate-specific contributions. In contrast, our target is the between-group difference in mean response rather than predictive performance, with both the predictor function and the covariate distribution differing between groups.

In this paper, we introduce a model-agnostic decomposition (MAD) for the difference in mean response between two groups. We formulate MAD as a cooperative switching game whose players are a model switch and individual covariate switches, organized into two *a priori* unions: a singleton model union and a covariate union containing all covariate switches. We then employ the Owen value (Owen, 1977) for a two-stage hierarchical allocation. In the first stage, the quotient game allocates the mean-response difference across the two unions, yielding the aggregate structure and composition effects, a two-factor Shapley decomposition (Shorrocks, 2013). In the second stage, the payoff of the covariate union, which is the aggregate composition effect, is further allocated across its covariate switches, yielding covariate-specific composition effects. While the first stage is determined entirely by the predictor functions and covariate distributions at the two endpoints, the second stage requires partial covariate switches, which create hybrid covariate vectors combining some covariates from one group with the remaining covariates from the other. The distribution of such a hybrid covariate vector is not determined by the group-specific covariate distributions alone. We therefore construct all hybrid covariate vectors from a single cross-group coupling, that is, a joint distribution pairing covariate vectors while preserving each group’s covariate distribution. To this end, we propose a minimum-switching-distortion (MSD) coupling, formulated as an optimal-transport problem (Villani, 2009).

MAD offers several advantages. First, it is model-agnostic, imposing no particular functional form and thereby accommodating modern machine-learning models that can flexibly adapt to complex nonlinear and interactive relationships. Second, its two-stage hierarchical allocation inherits the fair allocation properties of the Owen value, including exact two-level adding-up, so that the aggregate effects sum to the total difference and the covariate-specific composition effects sum to the aggregate composition effect. Third, its aggregate effects are invariant to the coupling, and under linear predictor functions, MAD reduces to the classical Blinder–Oaxaca decomposition with midpoint reference coefficients (Reimers, 1983). Finally, the MSD coupling is admissible, reducing to the identity coupling when the two covariate distributions coincide and remaining equivariant to group-label exchange and common covariate relabeling, and it has sparse support that reduces downstream allocation cost relative to alternative couplings.

We evaluate MAD through simulation studies, whose scenarios vary in the functional form of the covariate–response relationship, including nonlinearity and covariate interactions. For each scenario, we consider predictive models ranging from traditional linear regression to modern machine-learning models, namely gradient boosting machines (Friedman, 2001) and deep neural networks (LeCun et al., 2015), and assess the recovery of population structure and composition effects. The results show that MAD with machine-learning models yields more accurate decompositions than MAD with linear regression under nonlinear and interactive relationships, while linear regression remains competitive under linear additive relationships. We also compare the MSD coupling with two alternatives: independence and entropically regularized (Cuturi, 2013) couplings. The results show that, among the three couplings, only the MSD coupling satisfies identity at coincidence, so partial covariate switches leave the internal-game coalition value unchanged when the covariate distributions coincide. Moreover, its sparse support substantially reduces allocation cost relative to the alternatives.

We illustrate MAD using large-scale human microbiome data from the American Gut Project (McDonald et al., 2018) to decompose the difference in mean body mass index (BMI) between UK and US males aged 30–70 years into aggregate structure and composition effects and further allocate the aggregate composition effect across microbial genera. We discuss the interpretation of the genus-specific composition effects and their implications.

## 2 Methods

We first introduce notation and models, and review the classical Blinder–Oaxaca decomposition (Blinder, 1973; Oaxaca, 1973) under linear regression. We then formulate the model-agnostic decomposition (MAD) using a cooperative switching game and the Owen value (Owen, 1977), introduce the minimum-switching-distortion (MSD) coupling for hybrid covariate vectors, establish their theoretical properties, and describe computational procedures.

### 2.1 Notation and Models

Suppose that there are two groups, *A* and *B*. For each group *g* ∈ {*A, B*}, we let *Y*_*g*_ ∈ℝ denote a response with finite mean 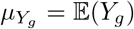 and finite variance 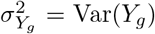, and let *X*_*g*_ = (*X*_*g*1_, …, *X*_*gp*_)^*⊤*^ ∈ ℝ^*p*^ denote a vector of *p* covariates with joint distribution *P*_*g*_, mean vector 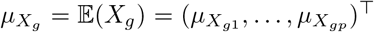, and covariance matrix 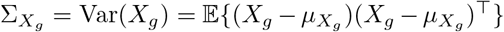, where 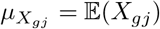 and 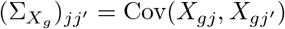 are finite for *j* = 1, …, *p* and *j*^*′*^ = 1, …, *p*.

We consider the model

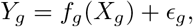

where *ϵ*_*g*_ is an error term satisfying E(*ϵ*_*g*_ | *X*_*g*_) = 0, and *f*_*g*_ may be nonlinear and may include covariate interactions.

The conditional and marginal means are then

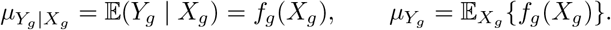

Finally, we define the total difference in mean response between the two groups as

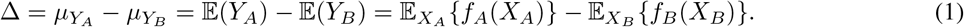

Two distinct sources contribute to Δ in Eq. (1): the groups may differ in the distributions of their covariates (*P*_*A*_≠ *P*_*B*_) and in their predictor functions (*f*_*A*_≠ *f*_*B*_). Disentangling these two sources is the object of what follows.

### 2.2 Blinder–Oaxaca Decomposition

The Blinder–Oaxaca decomposition (Blinder, 1973; Oaxaca, 1973) explains the total difference in mean response between the two groups under the linear regression model

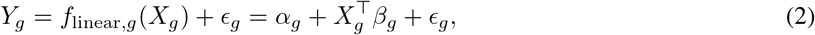

where *α*_*g*_ ∈ ℝ is the intercept and *β*_*g*_ = (*β*_*g*1_, …, *β*_*gp*_)^*⊤*^ ∈ℝ^*p*^ is the vector of slopes. Under Eq. (2), the marginal mean is given by

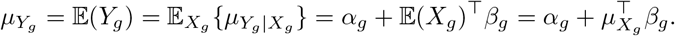

The total difference is then decomposed into structure and composition effects as

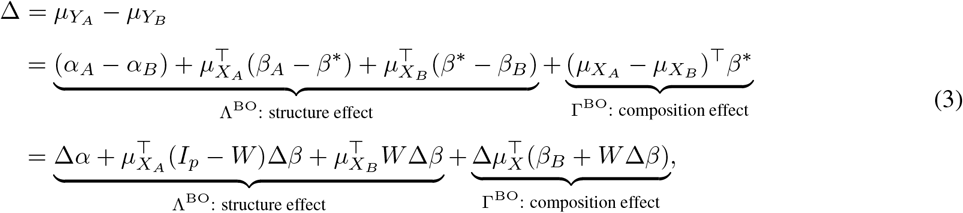

where 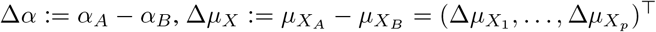, and Δ*β* := *β*_*A*_ − *β*_*B*_ = (Δ*β*_1_, …, Δ*β*_*p*_)^*⊤*^. Here, *β*^*\**^ is a reference coefficient vector of the form *β*^*\**^ = *Wβ*_*A*_ + (*I*_*p*_ − *W*)*β*_*B*_, where *W* = diag(*w*_1_, …, *w*_*p*_) with 0 ≤ *w*_*j*_ ≤ 1 for *j* = 1, …, *p*, and *I*_*p*_ denotes the *p* × *p* identity matrix. When 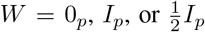, the reference coefficient vector *β*^*\**^ equals *β*_*B*_, *β*_*A*_, or 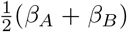, respectively (Blinder, 1973; Oaxaca, 1973; Reimers, 1983); alternatives include group-share weighting (Cotton, 1988) and pooled-regression coefficients (Neumark, 1988). Note that the structure effect Λ^BO^ reflects differences in the intercept and slopes, Δ*α* and Δ*β*, whereas the composition effect Γ^BO^ reflects differences in covariate means, Δ*µ*_*X*_, evaluated at *β*^*\**^.

The decomposition can further be expressed componentwise as

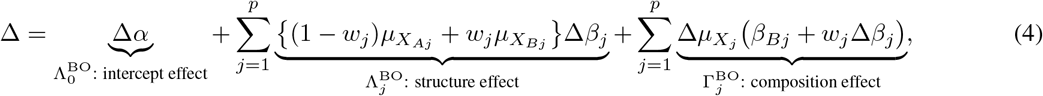

where 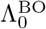 denotes the intercept channel, and 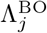 and 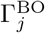 denote the structure and composition channels, respectively, for covariate *j*. This componentwise representation provides a complete additive decomposition across channels, 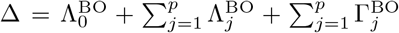, and satisfies channel-specific vanishing properties: 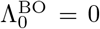 if Δ*α* = 0, 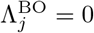 if Δ*β*_*j*_ = 0, and 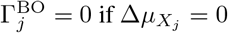. Each component is also linear in its corresponding difference term, holding the remaining quantities fixed.

### 2.3 Game-Theoretic Attribution with Minimum-Switching-Distortion Coupling

#### 2.3.1 Cooperative Switching Game

Throughout, we take group *B* as the reference group and group *A* as the comparison group, so that all switches proceed from group *B* to group *A*. Exchanging the group labels simply reverses the sign of each effect under an admissible coupling rule, as shown in Section 2.3.4.

##### Switches, players, and unions

The cooperative switching game is built on two types of switches: a model switch replaces *f*_*B*_ by *f*_*A*_, whereas a covariate switch replaces group-*B* covariates by their group-*A* counterparts. A covariate switch is full if all *p* covariates are switched and partial otherwise.

To formalize partial covariate switches, we index the covariates by [*p*] := {1, …, *p*}. For a subset *H* ⊆ [*p*] of switched covariates and its complement *H*^*c*^ := [*p*] \ *H* of retained covariates, we denote the corresponding subvectors of *X*_*g*_ = (*X*_*g*1_, …, *X*_*gp*_)^*⊤*^ by

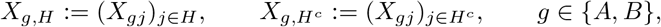

which have dimensions |*H*| and *p* − |*H*|, respectively. To encode the source group of each coordinate, we define

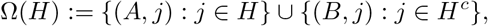

so that the coordinates in *H* are taken from group *A* and those in *H*^*c*^ are retained from group *B*. The corresponding hybrid covariate vector *X*_Ω(*H*)_ is then defined coordinatewise as

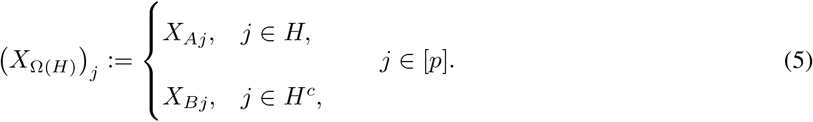

Thus, unlike the lower-dimensional subvectors *X*_*g,H*_ and 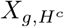, *X*_Ω(*H*)_ is a full *p*-dimensional vector that combines *X*_*A,H*_ on the switched coordinates with 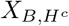 on the retained coordinates, in the original order of the covariates. At the two endpoints, Eq. (5) gives

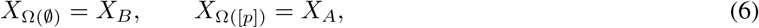

corresponding to no covariate switch and the full covariate switch, respectively.

We then treat the model switch and the *p* individual covariate switches as players, with

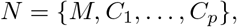

where *M* denotes the model switch and *C*_*j*_ the switch of covariate *j*. A coalition *S* ⊆ *N* specifies which switches are applied. To apply the Owen value (Owen, 1977), we partition *N* into two *a priori* unions,

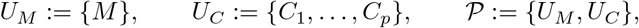

where *U*_*M*_ denotes the singleton model union, *U*_*C*_ denotes the covariate union, and *P* denotes the resulting partition.

##### Couplings for hybrid covariate vectors

At the two endpoints in Eq. (6), no coordinates from the two groups are combined: *X*_Ω(*∅*)_ = *X*_*B*_ ~ *P*_*B*_ and *X*_Ω([*p*])_ = *X*_*A*_ ~ *P*_*A*_, so both configurations are fully specified by the endpoint distributions *P*_*B*_ and *P*_*A*_ alone. For every intermediate *H* ⊆ [*p*], however, *X*_Ω(*H*)_ combines coordinates of *X*_*A*_ with coordinates of *X*_*B*_, which requires *X*_*A*_ and *X*_*B*_ to be defined on a common probability space. We therefore let *π* denote a coupling of *P*_*A*_ and *P*_*B*_, that is, a joint distribution on ℝ ^*p*^ ×ℝ ^*p*^ with marginals *P*_*A*_ and *P*_*B*_, respectively, and write

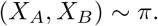

Thus, *π* specifies the cross-group joint distribution of *X*_*A*_ and *X*_*B*_, and hence the dependence between them. Given *π*, the hybrid covariate vector *X*_Ω(*H*)_ is well defined for every *H* ⊆ [*p*], and we denote its distribution by

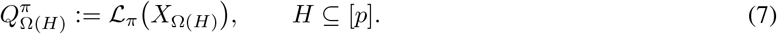

The *p* covariate switches yield 2^*p*^ hybrid covariate vectors. The coupling *π* is held fixed across coalitions, while the induced hybrid covariate distribution varies with *H*. The rule used to select *π* is specified in Section 2.3.3.

##### Characteristic function

For any coalition *S* ⊆ *N*, we define

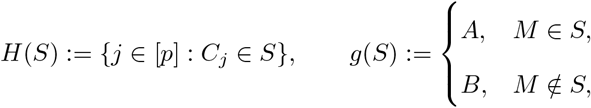

so that *H*(*S*) determines which covariates are switched, whereas *g*(*S*) determines which predictor function is used. Given *π*, we define the characteristic function *v*^*π*^ : 2^*N*^ →ℝ by

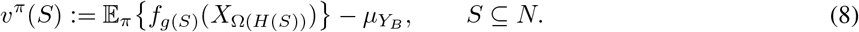

Thus, each coalition value is the change in the mean response relative to the baseline configuration of group *B*. At the two endpoints, Eqs. (6) and (8) give

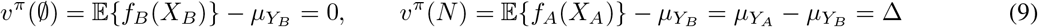

for any coupling *π*, corresponding to the empty coalition, which applies no switch, and the grand coalition, which applies all switches and returns the total difference Δ in Eq. (1), respectively.

#### 2.3.2 Two-Stage Hierarchical Allocation via the Owen Value

In our cooperative switching game (*N, v*^*π*^), the Owen value (Owen, 1977) first allocates Δ between the unions *U*_*M*_ and *U*_*C*_ via the quotient game and then allocates the payoff of *U*_*C*_ among *C*_1_, …, *C*_*p*_ via the internal game.

##### Stage 1: quotient game between the unions

In Stage 1, *U*_*M*_ and *U*_*C*_ are the two players of the quotient game associated with P (Owen, 1977), and the payoff of each union is its average marginal contribution over the two possible orders of the unions (Shapley, 1953). The marginal contributions involve only the four coalitions ∅, *U*_*M*_, *U*_*C*_, and *N*, in which either no covariate or all covariates are switched. Along with *v*^*π*^(∅) = 0 and *v*^*π*^(*N*) = Δ in Eq. (9), Eqs. (6) and (8) give

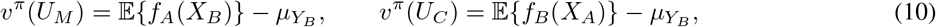

where *v*^*π*^(*U*_*M*_) switches the predictor function alone, whereas *v*^*π*^(*U*_*C*_) switches the covariate vector alone. We define the aggregate structure and composition effects as the payoffs of *U*_*M*_ and *U*_*C*_, respectively, which, by Eqs. (9) and (10), are

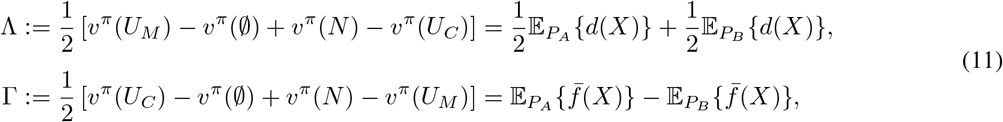

where *d*(*x*) := *f*_*A*_(*x*)−*f*_*B*_(*x*) and 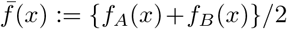 denote the difference and midpoint of the two predictor functions, respectively. The structure effect Λ averages the change from *f*_*B*_ to *f*_*A*_ under the two endpoint covariate distributions, whereas the composition effect Γ averages the change from *X*_*B*_ to *X*_*A*_ under the two group-specific predictor functions. Because both effects depend only on the endpoint predictor functions and covariate distributions, Stage 1 is coupling-invariant. By efficiency of the Shapley value (Shapley, 1953), the two union payoffs exhaust the total difference,

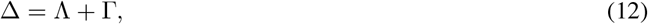

which is the two-factor Shapley decomposition of Δ in the sense of Shorrocks (2013).

##### Stage 2: internal game within the covariate union

In Stage 2, *C*_1_, …, *C*_*p*_ are the players of the internal game associated with *U*_*C*_, and the payoff of each covariate-switch player is its average marginal contribution over all possible orders of the covariate-switch players and the two possible orders of the unions (Owen, 1977). The marginal contributions involve the coalitions C_*H*_ := {*C*_*j*_ : *j* ∈ *H*} and *U*_*M*_ ∪ C_*H*_ for *H* ⊆ [*p*]. Given *π*, we define the covariate-specific composition effect of covariate *j* as the payoff of *C*_*j*_, which, by Eq. (8), is

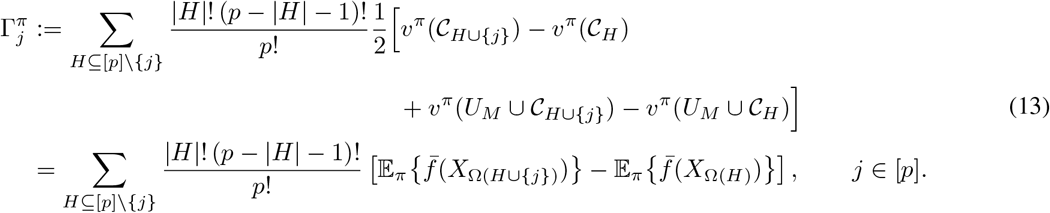

The second equality follows by averaging the two marginal contributions under *f*_*B*_ and *f*_*A*_, yielding 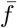. Thus, while Γ measures the effect of switching all covariates, 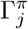 is the Shapley-weighted average marginal effect of switching covariate *j* over *H* ⊆ [*p*] \ {*j*}. Because 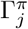 depends on the intermediate hybrid distributions induced by *π* in Eq. (7), Stage 2 is generally coupling-dependent. By the quotient-game property of the Owen value (Owen, 1977), for any coupling *π*, the payoffs of *C*_1_, …, *C*_*p*_ exhaust the payoff of *U*_*C*_,

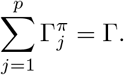

Thus, Stage 2 refines Γ into covariate-specific components without changing the Stage 1 split.

The construction above identifies Λ and 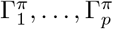 as the Owen payoffs of the model-switch and covariate-switch players, respectively. Beyond the two-level identities established above, the allocation inherits axioms for fair allocation from the Owen value, including null-player, within-union symmetry, and linearity (Owen, 1977). These results are summarized formally in the following proposition.

###### Proposition 1

(Owen representation and its consequences). *For any coupling π, the Owen allocation of* (*N, v*^*π*^) *under P assigns* Λ *to M and* 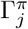, *and satisfies:*

i. Two-level coherence. *The covariate-specific composition effects exhaust the covariate-union payoff:*

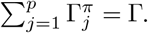
ii. *Efficiency. The structure effect and covariate-specific composition effects together exhaust the total difference:*

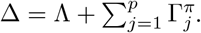
iii. Null player. *If v*^*π*^(*S* ∪ {*C*_*j*_}) = *v*^*π*^(*S*) *for every S* ⊆ *N* \ {*C*_*j*_}, *then* 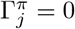.
iv. Symmetry within the covariate union. *If v*^*π*^(*S* ∪ {*C*_*j*_}) = *v*^*π*^(*S* ∪ {*C*_*k*_}) *for every S* ⊆ *N* \ {*C*_*j*_, *C*_*k*_}, *then* 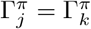.
v. Linearity. *If v*^*π*^ = *a*_1_*v*^*′*^ + *a*_2_*v*^*′′*^, *then each payoff is the corresponding linear combination of the Owen payoffs under v*^*′*^ *and v*^*′′*^ *with the same union structure P*.

##### The proof is given in Section S1 of the Supplementary Materials

The *a priori* union structure preserves the aggregate interpretation. Under a flat Shapley allocation on the full player set, intermediate hybrid covariate vectors can enter the payoff of *M*, so the model-switch payoff need not equal Λ and the covariate-switch payoffs need not sum to Γ. The hierarchical Owen allocation avoids this by preserving the quotient-game split and restricting the covariate-specific refinement to *U*_*C*_. A formal comparison is given in Section S2 of the Supplementary Materials.

Because Proposition 1 holds for any coupling *π*, only the internal-game allocation depends on the choice of *π* through the intermediate hybrid covariate vectors. We specify this choice next.

#### 2.3.3 Admissible Coupling Rules and the Minimum-Switching-Distortion Coupling Indeterminacy of the hybrid covariate distribution

By Eq. (5), *X*_Ω(*H*)_ combines the switched block *X*_*A,H*_ with the retained block 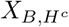, whose distributions are denoted by *P*_*A,H*_ and 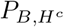, respectively. By the marginal constraints of the coupling, the hybrid covariate distribution 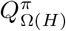 in Eq. (7) satisfies the block-marginal constraints

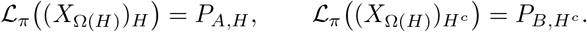

Thus, *P*_*A*_ and *P*_*B*_ determine the dependence within each block but not the cross-block dependence, because they do not specify how *X*_*A*_ is paired with *X*_*B*_. For an intermediate *H*, 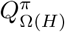is therefore generally not determined by *P*_*A*_ and *P*_*B*_ alone, and this cross-block dependence can affect 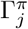 in Eq. (13) when 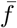 is nonadditive. We use a single coupling *π* to specify this cross-block dependence across all *H*, rather than a separate hybrid distribution for each *H*. Holding *π* fixed across coalitions ensures that each marginal contribution in Eq. (13) reflects only a covariate switch, not a simultaneous change in cross-group dependence. It remains to specify how *π* is selected.

##### Admissible coupling rules

Not every coupling rule is equally suitable for the decomposition. We call a coupling rule admissible if it is consistent at coincident endpoint distributions and equivariant to group exchange and common covariate relabeling, as formalized below.

###### Definition 1

(Admissible coupling rule). *A coupling rule on a domain D maps each* (*P*_*A*_, *P*_*B*_) ∈ *D to a coupling of P*_*A*_ *and P*_*B*_. *We call the rule admissible if it satisfies the following conditions*.

*(i)* Identity at coincidence. *If* (*P, P*) ∈ *D, the rule selects the identity coupling* L(*X, X*) *with X* ~ *P, so that X*_*A*_ = *X*_*B*_ *almost surely*.

*(ii)* Transposition equivariance. *If* (*P*_*A*_, *P*_*B*_) ∈ *D and* (*P*_*B*_, *P*_*A*_) ∈ *D, exchanging P*_*A*_ *and P*_*B*_ *transposes the selected coupling*.

*(iii)* Coordinate equivariance. *If a common relabeling of the covariates maps* (*P*_*A*_, *P*_*B*_) ∈ *D to another pair in D, the selected coupling is relabeled in the same way*.

Condition (i) selects the identity coupling when *P*_*A*_ = *P*_*B*_, so *X*_Ω(*H*)_ = *X*_*A*_ = *X*_*B*_ almost surely for every *H* ⊆ [*p*] and the internal-game coalition value is invariant to the covariate-switch subset. Conditions (ii) and (iii) ensure consistency under group exchange and common covariate relabeling, respectively. These conditions restrict the coupling rule without necessarily identifying a unique coupling.

##### MSD coupling

We select *π* by the MSD criterion, formulated as an optimal-transport problem that minimizes the expected squared standardized distance between paired *X*_*A*_ and *X*_*B*_. Specifically, we compare couplings by the expected switching cost

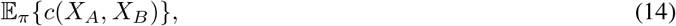

where

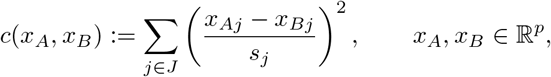

with

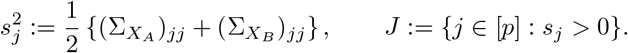

Here, *s*_*j*_ is a group-symmetric scale, and *J* indexes covariates that are nonconstant in at least one group. For *j* ∉ *J*, the covariate is constant within each group, so its paired difference is coupling-invariant; it is therefore excluded from the cost but remains a covariate-switch player. The symmetric scaling and coordinatewise sum make the criterion invariant to group exchange and common covariate relabeling. Standardization also makes it invariant to common coordinatewise affine transformations *x*_*j*_ → *a*_*j*_ + *b*_*j*_*x*_*j*_ with *b*_*j*_≠ 0. The criterion depends only on the covariate distributions, not on the response or predictor functions.

###### Definition 2

(MSD coupling). *We define the MSD coupling π*^*⋆*^ *as a minimizer of Eq*. (14) *over all couplings of P*_*A*_ *and P*_*B*_,

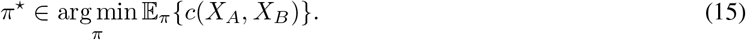

*If multiple minimizers exist, π^⋆^ is selected by a prespecified deterministic tie-breaking rule; the empirical implementation uses the maximum-entropy rule in Section 2.3.5*.

After standardizing the coordinates in *J* by *x*_*j*_ → *x*_*j*_*/s*_*j*_, Eq. (15) is the Kantorovich optimal-transport problem on ℝ^|*J*|^ with squared Euclidean cost (Villani, 2009; Peyré and Cuturi, 2019). Finite second moments ensure existence, while absolute continuity of either standardized endpoint distribution is a sufficient condition for uniqueness (Brenier, 1991; McCann, 1995). Details are given in Section S3 of the Supplementary Materials.

The invariances above yield conditions (ii) and (iii) of Definition 1. When *P*_*A*_ = *P*_*B*_, the identity coupling uniquely attains zero switching cost, yielding condition (i). Hence:

###### Proposition 2

(Admissibility). *On any domain where the MSD minimizer in Eq*. (15) *is unique, the coupling rule* (*P*_*A*_, *P*_*B*_) → *π*^*⋆*^ *is admissible. Under nonuniqueness, admissibility holds if the tie-breaking rule is equivariant to group exchange and common covariate relabeling. The proof is given in Section S3 of the Supplementary Materials*.

MAD uses the MSD coupling *π*^*⋆*^, with covariate-specific composition effects

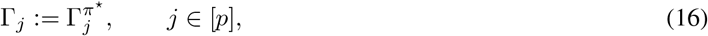

so that 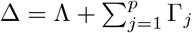 by Proposition 1.

##### Comparison with independence and entropically regularized couplings

The independence coupling *π*^ind^ := *P*_*A*_ ⊗ *P*_*B*_ pairs *X*_*A*_ and *X*_*B*_ independently, yielding 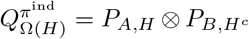 and thereby removing dependence between the switched and retained blocks. For *ε >* 0, the entropically regularized coupling is *π*^*ε*^:= arg min_*π*_ E_*π*_{*c*(*X*_*A*_, *X*_*B*_)} + *ε* KL(*π* ∥ *P*_*A*_ ⊗ *P*_*B*_), where KL denotes the Kullback–Leibler divergence. It approaches *π*^ind^ as *ε* → ∞ and, when the MSD minimizer is unique, *π*^*⋆*^ as *ε* → 0 (Cominetti and San Martín, 1994; Cuturi, 2013; Peyré and Cuturi, 2019). Both alternatives satisfy conditions (ii) and (iii) of Definition 1 but violate condition (i) whenever the common distribution is not a point mass (Section S4 of the Supplementary Materials). Thus, when *P*_*A*_ = *P*_*B*_, transposition equivariance still gives 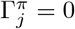 for every *j* ∈ [*p*], but partial covariate switches can change the internal-game coalition values when 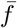 is nonadditive (Section 2.3.4).

#### 2.3.4 Further Properties and Connection to the Blinder–Oaxaca Decomposition

We establish further properties of MAD concerning coupling dependence, null cases, and group exchange.

##### Proposition 3

(Further properties of MAD). *MAD satisfies the following properties*.

i. Coupling invariance of the aggregate effects. *The aggregate effects* Λ *and* Γ *do not depend on the coupling*.
ii. Coupling invariance under an additive midpoint predictor. *If* 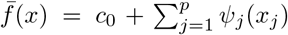, *where c*_0_ *is constant and the univariate functions ψ*_1_, …, *ψ*_*p*_ *have finite expectations under P*_*A*_ *and P*_*B*_, *then* 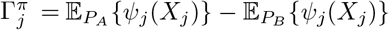 *for any coupling π and j* ∈ [*p*].
iii. Null structure effect. *If f*_*A*_(*x*) = *f*_*B*_(*x*) *for P*_*A*_*- and P*_*B*_*-almost every x, then* Λ = 0.
iv. Null composition effects at coincident distributions. *If P*_*A*_ = *P*_*B*_ = *P, then* Γ = 0 *for any coupling π. If the coupling is selected by a rule satisfying condition (ii) of Definition 1, then* 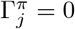 *for every j* ∈ [*p*]. *If the rule satisfies condition (i), then* 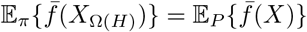 *for every H* ⊆ [*p*].
v. Group-exchange equivariance. *If the coupling is selected by a rule satisfying condition (ii) of Definition 1, exchanging groups A and B reverses the signs of* Λ, Γ, *and each covariate-specific composition effect*.

##### The proof is given in Section S5 of the Supplementary Materials

Parts (i) and (ii) characterize the role of the coupling: it does not affect Λ or Γ, and can affect the covariate-specific composition effects only when 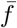 is nonadditive. Part (iv) separates two roles at coincidence: transposition equivariance makes the covariate-specific composition effects vanish, whereas identity at coincidence also makes the internal-game coalition value invariant to the covariate-switch subset. The MSD coupling satisfies both conditions by Proposition 2.

We next show that, under linear predictor functions, MAD reduces to the Blinder–Oaxaca decomposition with the midpoint reference coefficient vector of Reimers (1983).

###### Proposition 4

(Reduction to the Blinder–Oaxaca decomposition). *Suppose that f*_*g*_(*x*) = *α*_*g*_ + *x*^*⊤*^*β*_*g*_ *for g* ∈ {*A, B*}, *and let* Δ*α*, Δ*β*_*j*_, *and* 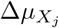 *be as in Eq*. (3). *Then, for any coupling π*,

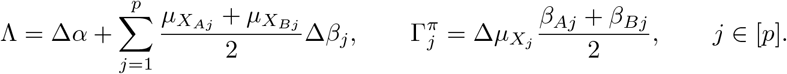

*Hence, with the components in Eq*. (4) *evaluated at* 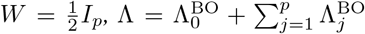 *and* 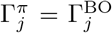 *for every j* ∈ [*p*]. *The proof is given in Section S6 of the Supplementary Materials*.

The reduction holds for any coupling *π* because linear predictor functions yield an additive midpoint predictor, so Proposition 3(ii) applies. Thus, MAD retains the classical decomposition as a special case, with the midpoint reference arising from the equal weighting of the two union orders, in line with the Shapley decomposition of Shorrocks (2013).

#### 2.3.5 Computational Procedures

For each group *g* ∈ {*A, B*}, we let {(*Y*_*gi*_, *X*_*gi*_) : *i* = 1, …, *n*_*g*_} be an independent random sample, and let 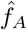 and 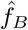 denote the fitted predictor functions. Treating the fitted predictor functions as given, we compute the plug-in cross-evaluated means as

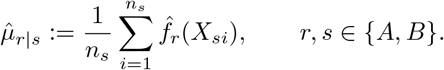

The total difference and aggregate effects are then

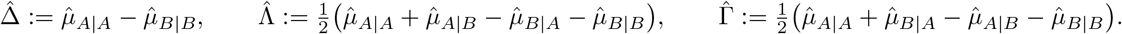

Thus, 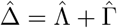 exactly, and these aggregate quantities require no coupling.

##### Empirical MSD coupling

We let 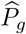 denote the empirical covariate distribution of group *g*, which assigns mass 1*/n*_*g*_ to each *X*_*gi*_, and define

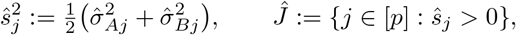

where 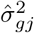 is the empirical variance computed with divisor *n*_*g*_. The pairwise switching costs are

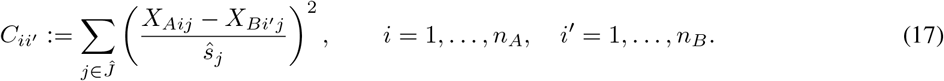

We let 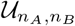 be the set of nonnegative *n*_*A*_ × *n*_*B*_ matrices 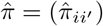 with row sums 1*/n*_*A*_ and column sums 1*/n*_*B*_, representing couplings of 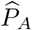 and 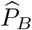. The empirical MSD problem with ground cost 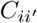 is

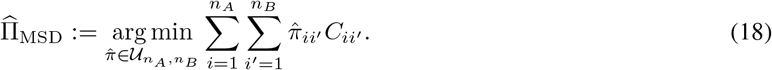

We select the maximum-entropy element of 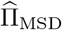, which is obtained as the *ε* → 0 limit of the corresponding entropically regularized transport problem (Cominetti and San Martín, 1994):

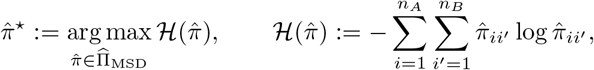

with 0 log 0 := 0. Strict concavity of H on the convex set 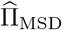 ensures uniqueness, and the resulting tie-breaking rule is equivariant as required by Proposition 2 (Section S3 of the Supplementary Materials).

##### Empirical independence and entropically regularized couplings

The empirical independence coupling 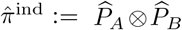 assigns 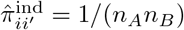 to every pair (*i, i*^*′*^) and requires no transport optimization or cost matrix. For *ε >* 0, the empirical entropically regularized coupling is 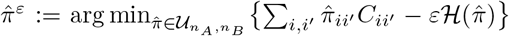, the empirical counterpart of 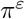, since 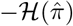 differs from KL 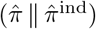 only by the constant log(*n*_*A*_*n*_*B*_) on 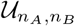. The coupling 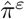 is unique and can be computed by Sinkhorn iterations (Cuturi, 2013; Peyré and Cuturi, 2019). Both couplings have full support on all *n*_*A*_*n*_*B*_ pairs.

##### Exact and approximate allocations

For each *H* ⊆ [*p*] and observation pair (*i, i*^*′*^), we define

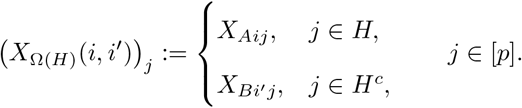

We let 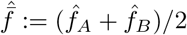 be the fitted midpoint predictor function and define

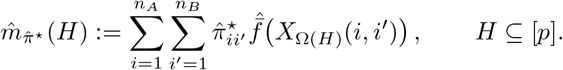

Since 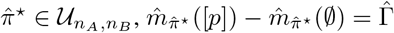. The exact covariate-specific allocation is

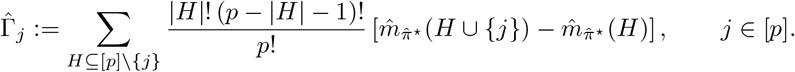

For larger *p*, the approximate allocation averages marginal contributions over sampled permutations (Castro et al., 2009; Mitchell et al., 2022). We let *ρ*_1_, …, *ρ*_*R*_ be independent uniform permutations of [*p*] and Pre_*j*_(*ρ*_*m*_) the set of covariates preceding *j* in *ρ*_*m*_, and we compute

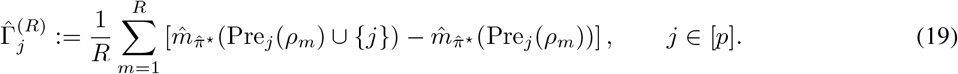

Since all covariates share the same permutations, the marginal contributions telescope within each permutation, so that 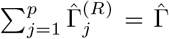 and 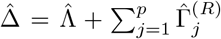 for every finite *R*. The budget *R* may be fixed or adaptive: pilot permutations determine a bounded budget targeting a prespecified relative Monte Carlo precision, and the allocation is computed from new independent permutations. For a fixed budget, and conditional on the pilot for an adaptive one, Eq. (19) is unbiased for the exact allocation (Section S7 of the Supplementary Materials). For 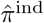 and 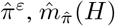 and the allocations are defined analogously, with 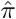 in place of 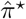.

##### Computational complexity

For computational comparison, suppose *n*_*A*_ = *n*_*B*_ = *n*. Then Eq. (18) reduces to a linear assignment problem (Kuhn, 1955), and under a unique optimal assignment the empirical MSD coupling has support size *n*. In contrast, the independence and entropically regularized couplings have support size *n*^2^, with the independence coupling requiring no transport optimization or cost-matrix construction. Table 1 summarizes the resulting leading costs. The smaller support of the empirical MSD coupling reduces the allocation cost by a factor of *n* relative to these alternatives, which lowers the overall cost whenever allocation dominates coupling construction. Additional details are given in Section S7 of the Supplementary Materials.

**Table 1.** Leading computational costs of the three empirical couplings under a unique optimal MSD assignment.

|  | MSD | Independence | Entropic |
|---|---|---|---|
| Coupling construction | $O(n^2p+n^3)$ | $O(1)$ | $O(n^2p+Tn^2)$ |
| Exact allocation | $O(2^pnC_{\mathrm{pred}}+p2^p)$ | $O(2^pn^2C_{\mathrm{pred}}+p2^p)$ | $O(2^pn^2C_{\mathrm{pred}}+p2^p)$ |
| Approximate allocation | $O(RpnC_{\mathrm{pred}})$ | $O(Rpn^2C_{\mathrm{pred}})$ | $O(Rpn^2C_{\mathrm{pred}})$ |

*Note.* $C_{\mathrm{pred}}$, cost of evaluating $\hat{\hat{f}}$ at one $p$-dimensional covariate vector; $T$, number of Sinkhorn iterations; $R$, number of sampled permutations.

## 3 Results

### 3.1 Simulation Designs

We conducted two simulation studies. First, the decomposition study evaluated MAD across predictive models and covariate–response structures, focusing on the recovery of population structure and composition effects and computational cost. Second, the coupling study compared the MSD coupling with the independence and entropically regularized couplings when the covariate distributions coincide, focusing on identity at coincidence, coupling sparsity, and computational cost.

#### Covariate dimensions and sample sizes

We considered *p* ∈ {5, 10, 15, 20} in both the decomposition and coupling studies, with *n*_*A*_ = *n*_*B*_ = *n* ∈ {100, 200} and an independent training sample of size *n*_train_ = 1000 per group. The corresponding coupling and allocation configurations are summarized in Table 2.

**Table 2.** Covariate dimensions, couplings, allocation procedures, and their roles.

| Study | Dimension | Coupling | Allocation | Role |
|---|---|---|---|---|
| Decomposition | $p \in \{5, 10, 15\}$ | MSD | Exact | Primary specification |
|  |  | MSD | Approximate | Allocation comparison |
|  | $p = 20$ | MSD | Approximate | Primary specification |
| Coupling | $p \in \{5, 10, 15\}$ | MSD, independence, entropic | Exact | Coupling comparison |
|  | $p = 20$ | MSD, independence, entropic | Approximate | Coupling comparison |

*Note*. Entropic: $\varepsilon = 0.05\ \mathrm{median}_{i,i'}(C_{ii'})$ with $C_{ii'}$ in Eq. (17); Approximate: fixed budget $R = 1000$ in Eq. (19).

#### Covariate distributions and predictor functions

We generated the covariate vectors as

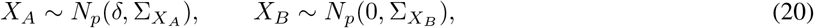

where *δ* is the between-group mean shift and 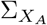 and 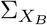 are the covariance matrices. We also considered four covariate–response structures: linear additive (LA), linear multiplicative (LM), nonlinear additive (NA), and nonlinear multiplicative (NM). Their true predictor functions are summarized in Table 3. The coefficient and distributional parameters in Eq. (20) and Table 3 are specified below.

**Table 3.** Four covariate–response relationship structures.

|  | Additive | Multiplicative |
|---|---|---|
| Linear | $\eta_A^{(\mathrm{LA})}(x) = \alpha + \sum_{j=1}^{p}(\beta_j + \gamma_j)x_j$<br>$\eta_B^{(\mathrm{LA})}(x) = \sum_{j=1}^{p} \beta_j x_j$ | $\eta_A^{(\mathrm{LM})}(x) = \alpha + \sum_{j=1}^{p}(\beta_j + \gamma_j)x_j + \sum_{k=1}^{K}(\theta_k + \xi_k)\chi_{A,k}(x)$<br>$\eta_B^{(\mathrm{LM})}(x) = \sum_{j=1}^{p} \beta_j x_j + \sum_{k=1}^{K} \theta_k \chi_{B,k}(x)$ |
| Nonlinear | $\eta_A^{(\mathrm{NA})}(x) = \alpha + \sum_{j=1}^{p}(\beta_j + \gamma_j)\phi_{r_A(j)}(x_j)$<br>$\eta_B^{(\mathrm{NA})}(x) = \sum_{j=1}^{p} \beta_j \phi_{r_B(j)}(x_j)$ | $\eta_A^{(\mathrm{NM})}(x) = \alpha + \sum_{j=1}^{p}(\beta_j + \gamma_j)\phi_{r_A(j)}(x_j) + \sum_{k=1}^{K}(\theta_k + \xi_k)\chi_{A,k}(x)$<br>$\eta_B^{(\mathrm{NM})}(x) = \sum_{j=1}^{p} \beta_j \phi_{r_B(j)}(x_j) + \sum_{k=1}^{K} \theta_k \chi_{B,k}(x)$ |

*Note. Interaction terms.* We used $K = 2p/5$ interaction terms, with two terms within each consecutive block of five covariates. For $q = 1, \ldots, p/5$, group $A$ uses $\chi_{A,2q-1}(x) = x_{5q-4}x_{5q-3}$ and $\chi_{A,2q}(x) = x_{5q-2}x_{5q-1}x_{5q}$; group $B$ reverses this assignment, with $\chi_{B,2q-1}(x) = x_{5q-2}x_{5q-1}x_{5q}$ and $\chi_{B,2q}(x) = x_{5q-4}x_{5q-3}$. *Nonlinear transformations.* We used $\phi_1(t) = \sin t$, $\phi_2(t) = \cos t - e^{-1/2}$, $\phi_3(t) = \tanh t$, and $\phi_4(t) = t^2 - 1$, each centered to have mean zero under $N(0,1)$. Group $A$ uses the forward cycle $r_A(j) = 1 + \{(j-1) \bmod 4\}$; group $B$ uses the reverse cycle $r_B(j) = 4 - \{(j-1) \bmod 4\}$.

In the decomposition study, we set

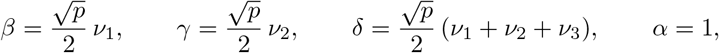

where *ν*_1_, *ν*_2_, *ν*_3_ were obtained by orthonormalizing three independent *N*_*p*_(0, *I*_*p*_) draws. Thus 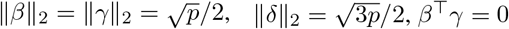, and *β*^*⊤*^*δ* = *γ*^*⊤*^*δ* = *p/*4. We let *K* = 2*p/*5, generated 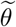 and 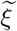 independently as normalized *N*_*K*_(0, *I*_*K*_) vectors, and set 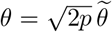 and 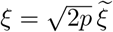, so the 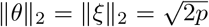. Finally, we specified 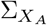 and 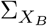 as independently generated unstructured correlation matrices by setting 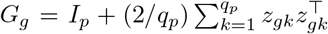 and 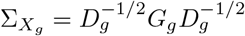, where 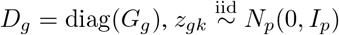, and *q* = ⌈*p/*2⌉, for *g* ∈ {*A, B*}.

In the coupling study, we set *δ* = 0 and 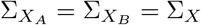, where Σ_*X*_ was generated using the same unstructured correlation construction, while retaining all other settings from the decomposition study, so that *P*_*A*_ = *P*_*B*_ = *P* = *N*_*p*_(0, Σ_*X*_), as in Proposition 3(iv). The same covariate sample was used for both groups with *X*_*A*_ = *X*_*B*_.

In both studies, the coefficient directions and unstructured correlation matrices were regenerated in each simulation replicate. In the decomposition study, the resulting *δ* was also replicate-specific, whereas the coupling study fixed *δ* = 0. Within each replicate, the realized coefficients and correlation matrices were shared across the two sample sizes and all covariate–response structures and, in the coupling study, across all coupling methods.

#### Response generation

For *g* ∈ {*A, B*} and *l* ∈ {LA, LM, NA, NM}, we generated

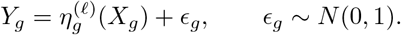

#### Predictive models

We considered five predictive models: linear regression; gradient boosting machines (Friedman, 2001), implemented using `XGBoost` (Chen and Guestrin, 2016) and `LightGBM` (Ke et al., 2017); and deep neural networks (LeCun et al., 2015), implemented using `Keras` (Chollet et al., 2015) and `Torch` (Paszke et al., 2019). Detailed training specifications are provided in Section S8 of the Supplementary Materials.

In addition, we considered an oracle benchmark using the true group-specific predictor functions, 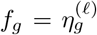. It removes model-estimation error, so the remaining error arises only from empirical covariate distributions and, under approximate allocation, permutation sampling. The coupling study used the oracle benchmark only.

#### Evaluation criteria

The evaluation criteria are summarized in Table 4. Each configuration was replicated *B* = 100 times.

**Table 4.**
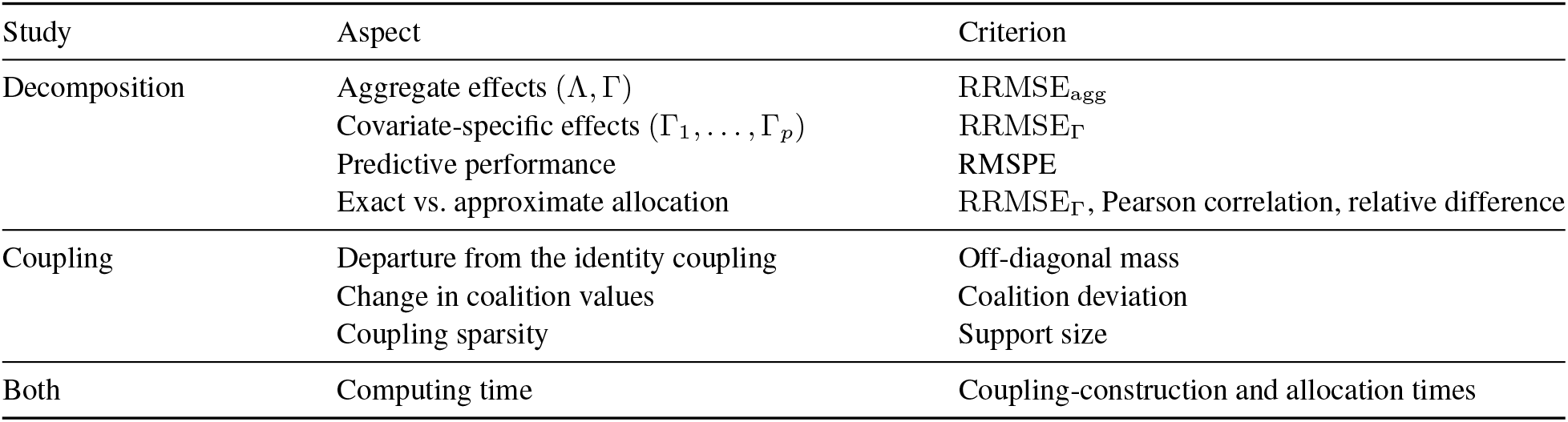
Evaluation criteria for the two simulation studies.

In the decomposition study, we measured the recovery of the aggregate and covariate-specific effects using the relative root mean squared errors (RRMSEs)

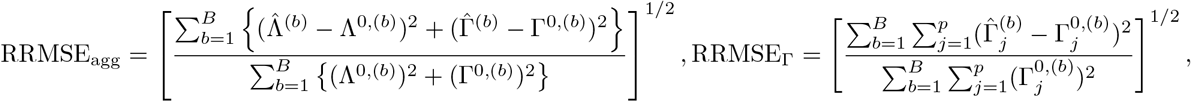

where *b* indexes simulation replicates. The population references Λ^0,(*b*)^, Γ^0,(*b*)^, and 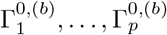 are described in Section S9 of the Supplementary Materials, with the covariate-specific references defined under the corresponding population MSD coupling. We also measured predictive performance using the root mean squared prediction error (RMSPE) of 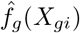 for *Y*_*gi*_ on the samples used for MAD, pooled over the two groups. Finally, we compared the exact and approximate allocations using RRMSE_Γ_ and assessed their agreement using the Pearson correlation and the relative root mean squared difference, 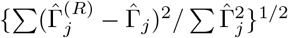, between paired 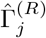 and 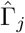 values pooled over replicates, covariates, and covariate–response structures.

In the coupling study, we measured three properties of each coupling 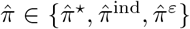: the departure from the identity coupling by the off-diagonal mass, 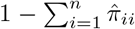; the change in coalition values by the coalition deviation, 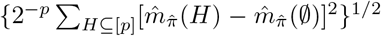, where 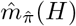 is the internal-game coalition value under 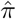, estimated from the sampled permutations for *p* = 20 (Section S10 of the Supplementary Materials); and the coupling sparsity by the support size, the number of pairs (*i, i*^*′*^) with 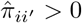.

In both studies, we recorded coupling-construction and allocation times.

### 3.2 Simulation Results

#### Decomposition study

Figure 1A shows the recovery of the aggregate and covariate-specific effects by MAD, measured by RRMSE_agg_ and RRMSE_Γ_, and the predictive performance, measured by RMSPE, for the five predictive models and the oracle benchmark. As expected, the oracle benchmark, which is correctly specified in every setting and thus free of model-estimation error, attained the smallest RRMSE_agg_, RRMSE_Γ_, and RMSPE throughout, that is, the most accurate decomposition and prediction. As *n* increased from 100 to 200, RRMSE_agg_ and RRMSE_Γ_ decreased consistently for the oracle benchmark, reflecting the smaller sampling error of the empirical covariate distributions.

**Figure 1.**
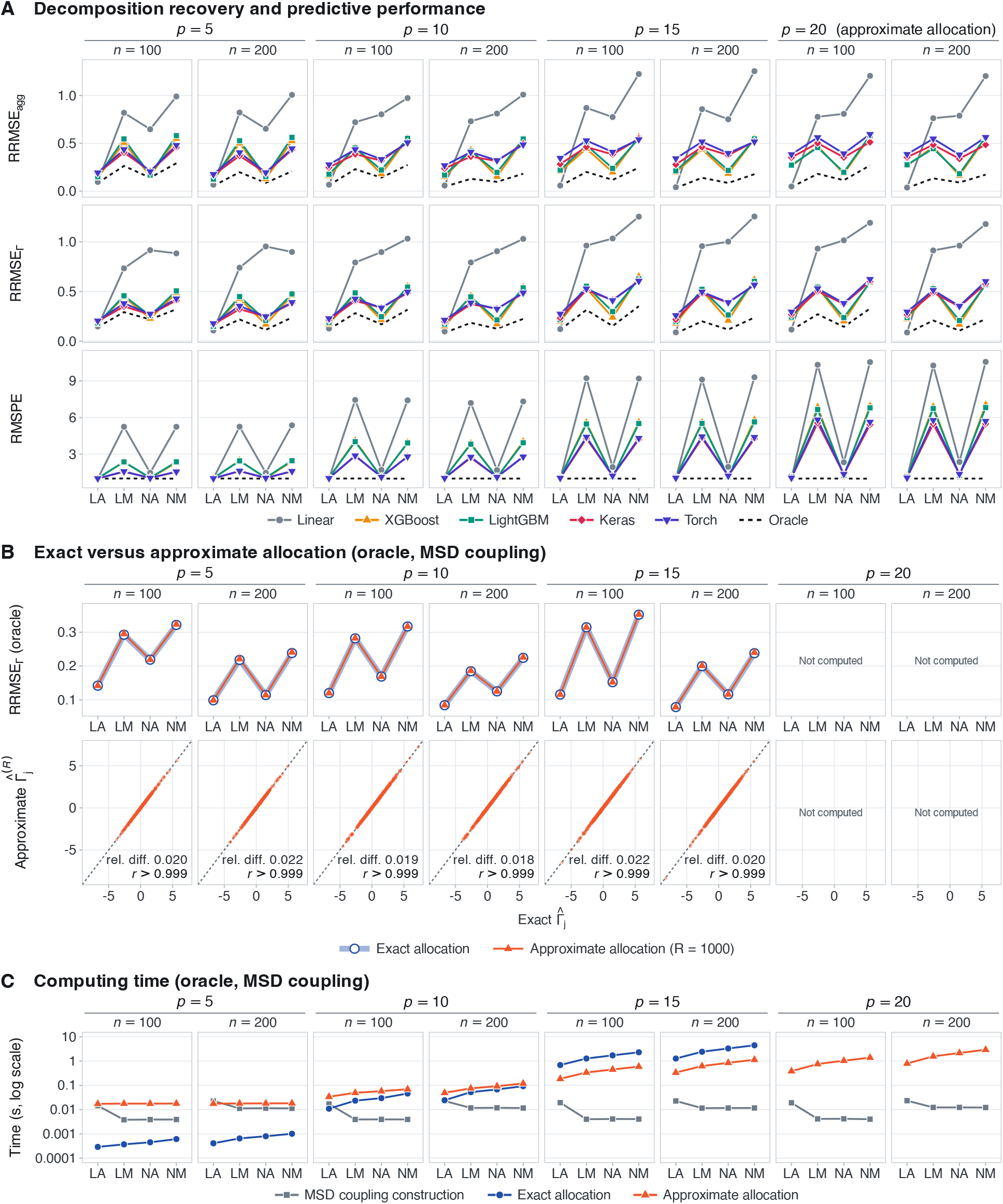
Decomposition study. (A) RRMSE_agg_, RRMSE_Γ_, and RMSPE of MAD with the five predictive models and the oracle benchmark under the MSD coupling, with the exact allocation for *p* ∈ {5, 10, 15} and the approximate allocation for *p* = 20. (B) RRMSE_Γ_ of the exact and approximate allocations (top) and approximate versus exact 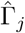 pooled over the four structures, with their relative root mean squared difference (rel. diff.) and Pearson correlation (*r*) (bottom), under the oracle benchmark. (C) Average computing times under the oracle benchmark.

Under the LA structure, where linear regression is correctly specified and MAD reduces to the midpoint Blinder– Oaxaca decomposition (Proposition 4), linear regression attained decomposition errors (RRMSE_agg_ and RRMSE_Γ_) close to those of the oracle benchmark, whereas the machine-learning models had larger decomposition errors that tended to increase with *p*. In contrast, under the LM, NA, and NM structures, which involve nonlinear transformations or covariate interactions, linear regression had large decomposition errors, whereas the machine-learning models substantially reduced them. The prediction error (RMSPE) showed the same contrast between linear regression and the machine-learning models, indicating that decomposition accuracy depended on how well the predictive model captured the covariate–response relationship. Thus, the flexibility of MAD in the choice of predictive model translated into more accurate decompositions under nonlinear and interactive relationships.

Figure 1B compares the exact and approximate allocations under the oracle benchmark for *p* ∈ {5, 10, 15}. Their RRMSE_Γ_ values differed by at most 0.003, and the paired 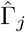 values had Pearson correlations above 0.999 and relative root mean squared differences of about 0.02. The approximate allocation, which is unbiased for the exact allocation and preserves exact adding-up (Section S7 of the Supplementary Materials), therefore closely reproduced the exact allocation.

Figure 1C shows the average computing times under the oracle benchmark. Constructing the MSD coupling took less than 0.03 seconds. The exact allocation time grew rapidly with *p*, reflecting the 2^*p*^ coalition values (Table 1), and reached 0.7–4.5 seconds for *p* = 15. The approximate allocation time, whose leading cost scales with *Rp* rather than 2^*p*^, grew more gradually, so that the approximate allocation was about four times faster than the exact allocation for *p* = 15 and took at most 3 seconds for *p* = 20, providing a scalable alternative for larger *p*.

#### Coupling study

Figure 2 summarizes the coupling study. As expected from Proposition 2, the empirical MSD coupling recovered the identity coupling, with zero off-diagonal mass in every replicate (Figure 2A). In contrast, the off-diagonal mass of the independence coupling was 1 − 1*/n*, and that of the entropically regularized coupling ranged from 0.004 to 0.34, decreasing with *p* and increasing with *n*, reflecting the violation of identity at coincidence by both alternatives (Section S4 of the Supplementary Materials). Accordingly, the coalition deviation of the MSD coupling remained at the round-off level, of the order of 10^*−*16^, whereas those of the independence and entropically regularized couplings averaged 0.21–0.32 and 3.3 × 10^*−*4^–1.3 × 10^*−*3^, respectively (Figure 2B). Hence, consistent with Proposition 3(iv), only the MSD coupling left the internal-game coalition values unchanged under partial covariate switches for every covariate–response structure when the covariate distributions coincided, whereas the two alternatives altered these values when the midpoint predictor was nonadditive (Proposition 3(ii)).

**Figure 2.**
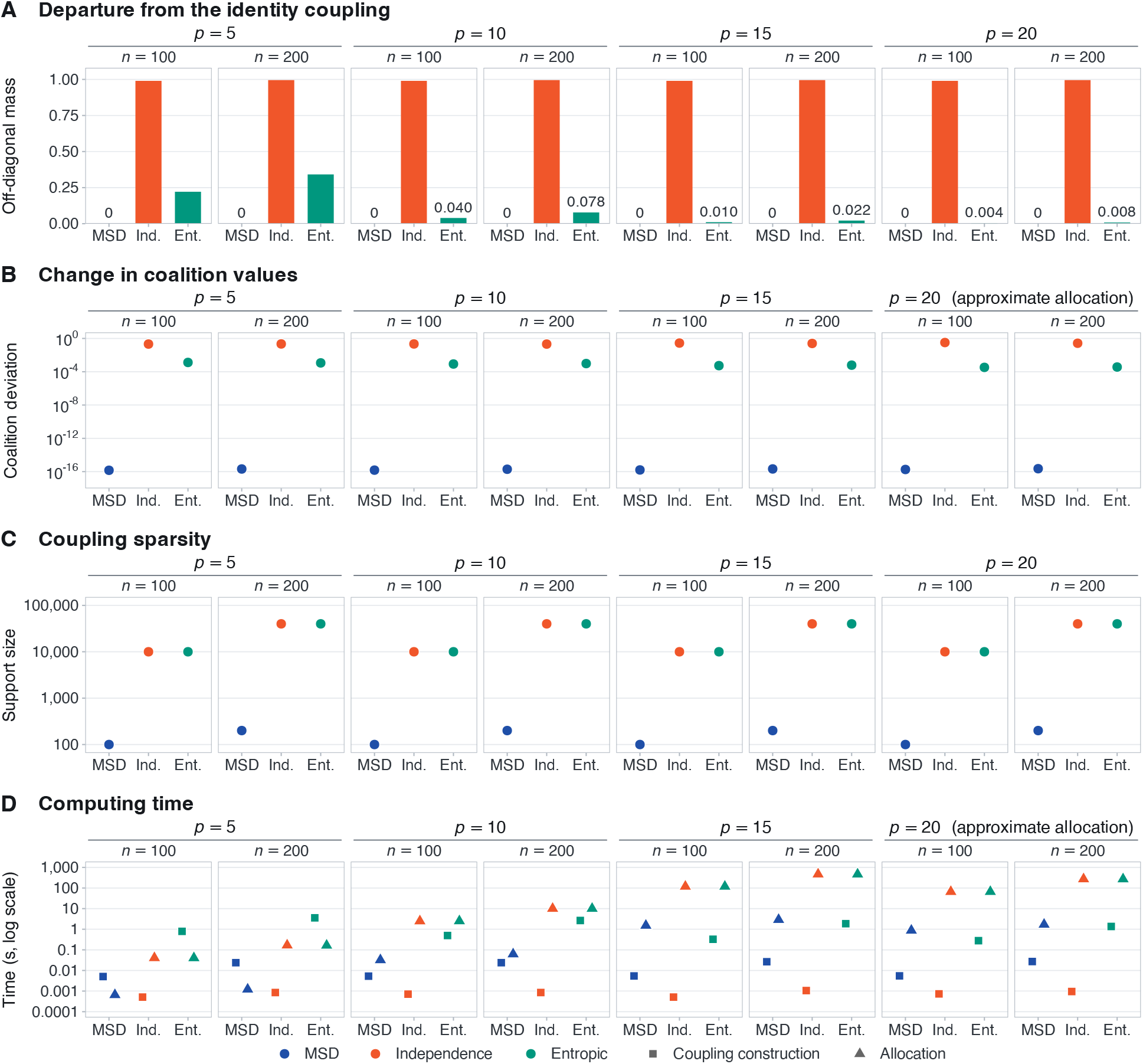
Coupling study under the oracle benchmark with *X*_*A*_ = *X*_*B*_. (A) Off-diagonal mass, (B) coalition deviation, (C) support size, and (D) average coupling-construction and allocation times of the MSD, independence (Ind.), and entropically regularized (Ent.) couplings, with the exact allocation for *p*∈{ 5, 10, 15} and the approximate allocation for *p* = 20. Coalition deviations and allocation times are averaged over the four covariate–response structures; for *p* = 20, coalition deviations were estimated from the sampled permutations (Section S10 of the Supplementary Materials).

The empirical MSD coupling had support size *n*, whereas the independence and entropically regularized couplings had full support of size *n*^2^ (Figure 2C). Constructing the MSD coupling took less than 0.03 seconds, compared with 0.3–3.6 seconds for the entropically regularized coupling, whereas the independence coupling required no optimization (Figure 2D). Owing to its sparse support, the MSD coupling reduced the allocation time relative to the other two couplings by about two orders of magnitude, with factors of 75–78 for *n* = 100 and about 160 for *n* = 200 when *p* ≥ 10, consistent with the factor-of-*n* reduction in Table 1. For *p* = 15 and *n* = 200, for example, the allocation took 2.9 seconds under the MSD coupling and about 470 seconds under the other two couplings. Overall, the MSD coupling was the only one of the three couplings to satisfy identity at coincidence, and its sparse support also lowered allocation cost.

## 4 Real Data Applications

We illustrate the proposed MAD with the MSD coupling using 16S rRNA gene amplicon data from the American Gut Project (AGP) (McDonald et al., 2018). In particular, we decompose the difference in mean body mass index (BMI, kg/m^2^) between UK and US males aged 30–70 years, with the UK as the comparison group *A* and the US as the baseline group *B*. We applied data quality control and transformation procedures as follows. First, we retained one fecal sample per participant with at least 10,000 reads and a BMI of at least 15 and below 60, yielding *n*_*A*_ = 730 UK and *n*_*B*_ = 1,864 US males, and the microbial genera with a mean relative abundance of at least 0.001 among all eligible participants, yielding *p* = 53 genera. Second, we transformed the genus counts into centered log-ratio (CLR) coordinates (Aitchison, 1986), with zero counts replaced by 0.5, which map the compositions isometrically from the simplex onto a hyperplane of the Euclidean space (Egozcue et al., 2003). The mean BMIs of the UK and US males were 24.91 and 25.44, respectively, giving an observed difference of −0.53, which was significant at the 0.05 level by the Welch *t*-test (*P* = 0.003; Welch, 1947).

For each country, BMI was predicted from the CLR coordinates of the *p* = 53 genera using linear regression, gradient boosting machines (`XGBoost` and `LightGBM`), and deep neural networks (`Keras` and `Torch`). `LightGBM` was selected as the predictive model because it had the smallest cross-validated root mean squared error. Because a hybrid covariate vector that combines the CLR coordinates of a UK and a US participant no longer lies on the CLR hyperplane, each predictor was evaluated at the orthogonal projection of the hybrid covariate vector onto this hyperplane, which leaves the predictions for all observed participants unchanged. We then applied the MSD coupling and the adaptive-budget allocation (Section 2.3.5) with a tolerance of 0.0045, slightly below the target relative Monte Carlo standard error of 0.005, which required *R* = 646 permutations and achieved 0.0045. Data processing, training procedures, predictive performance, and decomposition settings are detailed in Section S11 of the Supplementary Materials.

Figure 3A shows that, of the total difference Δ = −0.528, the aggregate structure effect was Λ = −0.428 (81%) and the aggregate composition effect was Γ = −0.100 (19%). A substantial share of the lower mean BMI of UK males is thus attributed to differences between the two countries in how the genus composition relates to BMI, rather than to differences in the genus composition itself. Such structural differences may reflect how the host interacts with and regulates its gut microbiota, which can be governed by host factors such as genetics and physiology. Since the predictors include the genus composition alone, however, Λ also absorbs between-country differences in other determinants of BMI that were not included as covariates, such as age, diet, living environment, and physical activity.

**Figure 3.**
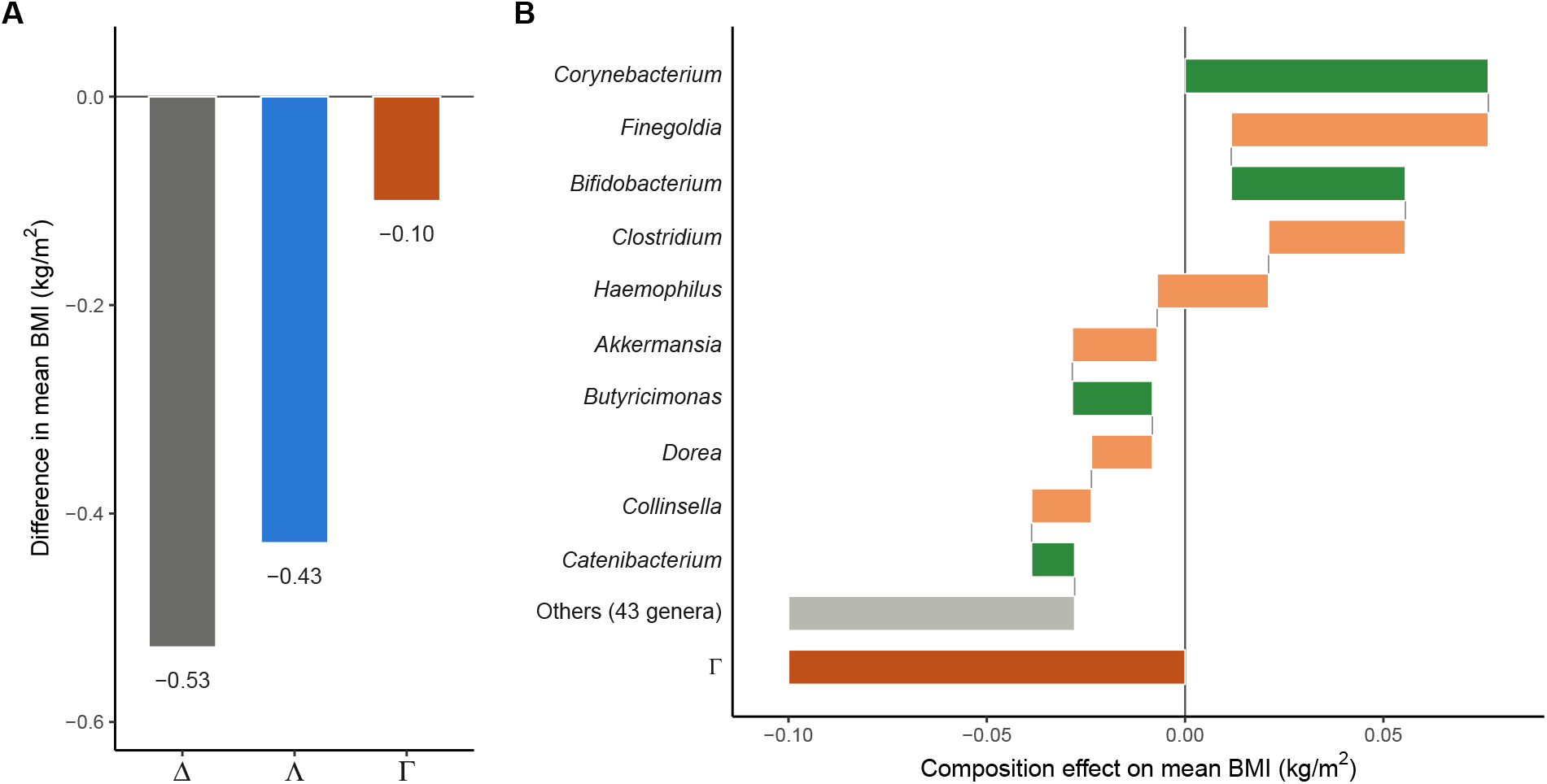
Decomposition of the difference in mean BMI (kg/m^2^) between UK (*n*_*A*_ = 730) and US (*n*_*B*_ = 1,864) males aged 30–70 years in the American Gut Project, based on `LightGBM` from the CLR coordinates of 53 genera, the MSD coupling, and the adaptive-budget allocation. (A) Total difference Δ (gray), aggregate structure effect Λ (blue), and aggregate composition effect Γ (dark orange), with Δ = Λ + Γ. (B) Waterfall of the genus-specific composition effects Γ_*j*_ of the top ten genera, followed by the sum over the remaining 43 genera (Others, gray) and Γ; positive and negative effects are shown in green and light orange, respectively. All effects are UK minus US.

Figure 3B shows the genus-specific composition effects Γ_*j*_ as a waterfall chart, in which the bars cumulatively add the effects of the top ten genera with the largest |Γ_*j*_| and then the combined effect of the remaining 43 genera to reach Γ; green and orange bars indicate composition differences associated with a higher and a lower mean BMI in the UK, respectively. The top ten genera contributed only −0.028 because their positive and negative effects largely offset each other, whereas the remaining 43 genera contributed −0.072. The two largest effects, those of *Corynebacterium* (0.076) and *Finegoldia* (−0.065), nearly offset each other. *Bifidobacterium*, a probiotic genus familiar from yogurt and other fermented dairy products, has been inversely associated with obesity (Million et al., 2012), and *Akkermansia*, a mucus-degrading genus that helps maintain the intestinal barrier, has been shown experimentally to counteract diet-induced obesity and metabolic inflammation (Everard et al., 2013). In mean relative abundance, *Bifidobacterium* was slightly less abundant in the UK than in the US (1.47% vs. 1.58%), whereas *Akkermansia* was more abundant in the UK than in the US (2.44% vs. 2.08%). Their composition effects had opposite signs: the positive effect of *Bifidobacterium* (Γ_*j*_ = 0.044) offset part of the lower mean BMI of UK males, whereas the negative effect of *Akkermansia* (Γ_*j*_ = −0.021) added to it, consistent with previous findings. These genera may therefore merit investigation as targets of dietary, probiotic, or prebiotic interventions, though the substantial structure effect suggests that such interventions may need to be complemented by attention to other determinants of BMI.

As the AGP is observational, these effects are descriptive rather than causal. On a single CPU thread, the MSD coupling took 0.5 seconds, and the adaptive-budget allocation took 18.0 minutes.

## 5 Discussion

In this paper, we proposed MAD, a model-agnostic decomposition of the difference in mean response between two groups into structure and composition effects. MAD formulates a cooperative switching game with a model-switch player and individual covariate-switch players, organized into model and covariate unions. It then employs the Owen value (Owen, 1977) for a two-stage hierarchical allocation, in which the quotient game yields the aggregate structure and composition effects and the internal game allocates the aggregate composition effect across covariates. MAD thereby inherits the fair allocation properties of the Owen value, including exact two-level adding-up, and reduces to the midpoint Blinder–Oaxaca decomposition under linear predictor functions. For the hybrid covariate vectors arising in the internal game, the MSD coupling satisfies identity at coincidence, transposition equivariance, and coordinate equivariance, and has sparse support. The exact allocation is feasible for moderate *p*, and the approximate allocation based on sampled permutations provides a scalable alternative that preserves exact adding-up.

The simulation studies supported the accuracy of MAD and the advantages of the MSD coupling. MAD with machine-learning models recovered the population structure and composition effects more accurately than MAD with linear regression under nonlinear and interactive relationships, while linear regression remained competitive under linear additive relationships, and the approximate allocation closely agreed with the exact allocation. Among the three couplings, only the MSD coupling satisfied identity at coincidence, and its sparse support substantially reduced allocation cost. The application to the American Gut Project (McDonald et al., 2018) further illustrated the practical utility of MAD: the lower mean BMI of UK males relative to US males was attributed largely to the aggregate structure effect (81%) and partly to the aggregate composition effect (19%), with the genus-specific effects of *Bifidobacterium* and *Akkermansia* consistent with previous findings. Together, these results suggest that MAD can provide accurate and interpretable decompositions with modern machine-learning models.

As with other attribution methods based on fitted models, the effects estimated by MAD depend on how well the predictive models capture the covariate–response relationships, and, with observational data, they are descriptive rather than causal. Several extensions merit future research. First, MAD could be extended toward causal inference by combining it with appropriate identification assumptions and causal estimation methods, so that the structure and composition effects admit causal interpretations. Second, statistical inference for the estimated effects, accounting for both model estimation and sampling variability, would complement the point estimates. Third, MAD could be extended to other outcome structures, such as survival outcomes and longitudinal or time-series data, with suitable target quantities such as survival probabilities and time-indexed mean trajectories.

Overall, MAD provides a principled and computationally practical framework for decomposing between-group differences in mean response with modern machine-learning models. Although we illustrated MAD with microbiome data, its model-agnostic formulation can be readily applied to other high-dimensional omics data, such as genomic, proteomic, and metabolomic data, and to between-group comparisons beyond biomedical research, such as in engineering and the social sciences, particularly where covariate–response relationships are complex.

## Supplementary Materials

Sections S0–S7 of the Supplementary Materials provide proofs of Propositions 1–4, properties of alternative couplings, and computational details of approximate allocation. Sections S8–S10 describe the training specifications, population reference quantities, and coalition deviation estimation under approximate allocation for the simulation studies. Section S11 details the real data application, including the genus-specific composition effects for all 53 genera.

## Data Availability

This study is a secondary analysis of publicly available, de-identified 16S rRNA gene amplicon data from the American Gut Project (AGP) (McDonald et al., 2018). The sequence data, sample metadata, and processed feature tables are freely available from Qiita (https://qiita.ucsd.edu; study ID 10317).

## Funding

This work was supported by the National Research Foundation of Korea (NRF) grant funded by the Korean government (MSIT) (No. RS-2026-25477838).

## Competing Interests

The author declares no competing interests.

## Notes

### Competing Interest Statement

The authors have declared no competing interest.

