## Supplementary Materials for "Model-Agnostic Decomposition of the Difference in Mean Response Between Two Groups via Game-Theoretic Attribution with Minimum-Switching-Distortion Coupling"

**Hyunwook Koh**

**Department of Applied Mathematics and Statistics**
**The State University of New York, Korea**
**119 Songdo Moonhwa-ro, Yeonsu-gu, Incheon 21985, South Korea**

****

### Overview

This supplement provides proofs and technical details for MAD. Sections S0–S7 accompany the Methods section, covering the notation, the Owen representation, the comparison with a flat Shapley allocation, the MSD coupling and alternative couplings, further properties, the Blinder–Oaxaca reduction, and approximate allocation and computational complexity. Sections S8–S10 give the simulation training specifications, population reference quantities, and coalition-deviation calculation, and Section S11 gives the details of the real data application.

### S0. Notation

We follow the notation of the main text and collect here the quantities used in the proofs. We let $[p] := \{1, \ldots, p\}$ and, for $H \subseteq [p]$, $H^c := [p] \setminus H$. For group $g \in \{A, B\}$, we let $P_g$ denote the distribution of $X_g \in \mathbb{R}^p$, with mean vector $\mu_{X_g}$ and covariance matrix $\Sigma_{X_g}$, and $P_{g,H}$ the distribution of the subvector $X_{g,H}$, with $P_{g,j} := P_{g,\{j\}}$. We let $f_g : \mathbb{R}^p \to \mathbb{R}$ denote the group-specific predictor function, $f_g(x) = \mathbb{E}(Y_g \mid X_g = x)$, so that $\mu_{Y_g} = \mathbb{E}\{f_g(X_g)\}$ and $\Delta = \mu_{Y_A} - \mu_{Y_B}$. All expectations used below are assumed finite.

For $H \subseteq [p]$, the hybrid covariate vector is

$$\left(X_{\Omega(H)}\right)_j := \begin{cases} X_{Aj}, & j \in H, \\ X_{Bj}, & j \in H^c, \end{cases} \qquad j \in [p],$$

so that $X_{\Omega(\emptyset)} = X_B$ and $X_{\Omega([p])} = X_A$. A coupling $\pi$ of $P_A$ and $P_B$ is a joint distribution of $(X_A, X_B)$ on $\mathbb{R}^p \times \mathbb{R}^p$ with marginals $P_A$ and $P_B$, respectively, and $Q^{\pi}_{\Omega(H)} := \mathcal{L}_{\pi}(X_{\Omega(H)})$ denotes the distribution of the hybrid covariate vector under $\pi$. The marginal constraints of the coupling give

$$\mathcal{L}_{\pi}\left((X_{\Omega(H)})_H\right) = P_{A,H}, \qquad \mathcal{L}_{\pi}\left((X_{\Omega(H)})_{H^c}\right) = P_{B,H^c},$$

and, at the two endpoints,

$$Q^{\pi}_{\Omega(\emptyset)} = P_B, \qquad Q^{\pi}_{\Omega([p])} = P_A \tag{S1}$$

for any coupling $\pi$.

The cooperative switching game has player set and *a priori* union structure

$$N = \{M, C_1, \ldots, C_p\}, \qquad U_M := \{M\}, \qquad U_C := \{C_1, \ldots, C_p\}, \qquad \mathcal{P} := \{U_M, U_C\}.$$

For $S \subseteq N$, we let $H(S) := \{j \in [p] : C_j \in S\}$, and $g(S) := A$ if $M \in S$ and $g(S) := B$ otherwise. Given $\pi$, the characteristic function is

$$v^{\pi}(S) := \mathbb{E}_{\pi}\left\{f_{g(S)}(X_{\Omega(H(S))})\right\} - \mu_{Y_B}, \qquad S \subseteq N.$$

By Eq. (S1),

$$v^{\pi}(\emptyset) = 0, \quad v^{\pi}(U_M) = \mathbb{E}\{f_A(X_B)\} - \mu_{Y_B}, \quad v^{\pi}(U_C) = \mathbb{E}\{f_B(X_A)\} - \mu_{Y_B}, \quad v^{\pi}(N) = \Delta \tag{S2}$$

for any coupling $\pi$. With

$$d(x) := f_A(x) - f_B(x), \qquad \bar{f}(x) := \tfrac{1}{2}\{f_A(x) + f_B(x)\},$$

the aggregate structure and composition effects, the payoffs of $U_M$ and $U_C$ in the quotient game, are

$$\Lambda = \tfrac{1}{2}\mathbb{E}_{P_A}\{d(X)\} + \tfrac{1}{2}\mathbb{E}_{P_B}\{d(X)\}, \qquad \Gamma = \mathbb{E}_{P_A}\{\bar{f}(X)\} - \mathbb{E}_{P_B}\{\bar{f}(X)\}, \tag{S3}$$

with $\Lambda + \Gamma = \Delta$. Neither quantity depends on $\pi$.

For $H \subseteq [p]$, we let $\mathcal{C}_H := \{C_j : j \in H\}$, and we abbreviate the Shapley weights as

$$\kappa_k := \frac{k!\,(p-k-1)!}{p!}, \qquad k = 0, \ldots, p-1.$$

Given $\pi$, the covariate-specific composition effect of covariate $j$ is

$$\begin{aligned} \Gamma^{\pi}_j &= \sum_{H \subseteq [p]\setminus\{j\}} \kappa_{|H|} \frac{1}{2}\Big[v^{\pi}(\mathcal{C}_{H\cup\{j\}}) - v^{\pi}(\mathcal{C}_H) \\ &\qquad\qquad + v^{\pi}(U_M \cup \mathcal{C}_{H\cup\{j\}}) - v^{\pi}(U_M \cup \mathcal{C}_H)\Big] \\ &= \sum_{H \subseteq [p]\setminus\{j\}} \kappa_{|H|} \left[\mathbb{E}_{\pi}\{\bar{f}(X_{\Omega(H\cup\{j\})})\} - \mathbb{E}_{\pi}\{\bar{f}(X_{\Omega(H)})\}\right]. \end{aligned} \tag{S4}$$

MAD uses the MSD coupling $\pi^{\star}$ of Section S3, with $\Gamma_j := \Gamma^{\pi^{\star}}_j$.

### S1. Proof of the Owen Representation and Its Consequences

For a game $(N, v)$ with *a priori* unions $\mathcal{P} = \{U_1, \ldots, U_m\}$ and $i \in U_k$, we let $n_k := |U_k|$ and $U_R := \bigcup_{\ell \in R} U_\ell$ for $R \subseteq \{1, \ldots, m\}$. The Owen value (Owen, 1977) is

$$\mathrm{Ow}_i(N, v, \mathcal{P}) = \sum_{R \subseteq \{1,\ldots,m\} \setminus \{k\}} \sum_{T \subseteq U_k \setminus \{i\}} \frac{|R|!\,(m - |R| - 1)!}{m!} \frac{|T|!\,(n_k - |T| - 1)!}{n_k!} \{v(U_R \cup T \cup \{i\}) - v(U_R \cup T)\}. \qquad \text{[S5]}$$

**Proposition 1 (Owen representation and its consequences).** *For any coupling $\pi$, the Owen allocation of $(N, v^\pi)$ under $\mathcal{P}$ assigns $\Lambda$ to $M$ and $\Gamma_j^\pi$ to $C_j$, for $j \in [p]$, and satisfies: (i) two-level coherence, $\sum_{j=1}^{p} \Gamma_j^\pi = \Gamma$; (ii) efficiency, $\Delta = \Lambda + \sum_{j=1}^{p} \Gamma_j^\pi$; (iii) the null-player property: if $v^\pi(S \cup \{C_j\}) = v^\pi(S)$ for every $S \subseteq N \setminus \{C_j\}$, then $\Gamma_j^\pi = 0$; (iv) symmetry within the covariate union: if $v^\pi(S \cup \{C_j\}) = v^\pi(S \cup \{C_k\})$ for every $S \subseteq N \setminus \{C_j, C_k\}$, then $\Gamma_j^\pi = \Gamma_k^\pi$; and (v) linearity: if $v^\pi = a_1 v' + a_2 v''$, then each payoff is the corresponding linear combination of the Owen payoffs under $v'$ and $v''$ with the same union structure $\mathcal{P}$.*

*Proof.* Because $\mathcal{P}$ contains two unions and $U_M$ is a singleton,

$$\mathrm{Ow}_M = \tfrac{1}{2}\{v^\pi(U_M) - v^\pi(\emptyset)\} + \tfrac{1}{2}\{v^\pi(N) - v^\pi(U_C)\} = \Lambda$$

by Eq. (S2)–Eq. (S3). The quotient-game payoff of $U_C$ is similarly $\Gamma$, and Shapley efficiency gives $\Lambda + \Gamma = \Delta$.

For $C_j$, the two possible orders of the unions each have weight $1/2$, while each coalition $\mathcal{C}_H$ of covariate-switch players preceding $C_j$ within $U_C$ has weight $\kappa_{|H|}$. Substituting these weights into Eq. (S5) gives the first equality in Eq. (S4), so $\mathrm{Ow}_{C_j} = \Gamma_j^\pi$. Because $v^\pi(\mathcal{C}_H)$ and $v^\pi(U_M \cup \mathcal{C}_H)$ evaluate $f_B$ and $f_A$, respectively, at $X_{\Omega(H)}$, averaging the two marginal contributions for each $H$ gives the second equality in Eq. (S4). This is the ordinary Shapley formula (Shapley, 1953) for the set function

$$H \longmapsto \mathbb{E}_\pi\{\bar{f}(X_{\Omega(H)})\}.$$

Therefore, by Shapley efficiency and Eq. (S1),

$$\sum_{j=1}^{p} \Gamma_j^\pi = \mathbb{E}_{P_A}\{\bar{f}(X)\} - \mathbb{E}_{P_B}\{\bar{f}(X)\} = \Gamma. \qquad \text{[S6]}$$

Equation Eq. (S6) proves two-level coherence, and combining it with $\Lambda + \Gamma = \Delta$ proves efficiency.

The null-player, within-union symmetry, and linearity properties follow directly from the corresponding properties of the Owen value, or equivalently from Eq. (S5). ☐

### S2. Formal Comparison with a Flat Shapley Allocation

Given $\pi$, we define

$$\Lambda^{\text{flat}}(\pi) := \mathrm{Sh}_M(N, v^\pi), \qquad \Gamma_j^{\text{flat}}(\pi) := \mathrm{Sh}_{C_j}(N, v^\pi)$$

as the Shapley payoffs (Shapley, 1953) under a flat allocation on the full player set $N$ that ignores the *a priori* union structure. The model-switch payoff is

$$\Lambda^{\text{flat}}(\pi) = \sum_{H \subseteq [p]} \omega_{|H|} \mathbb{E}_\pi\{d(X_{\Omega(H)})\}, \qquad \omega_k := \frac{k!\,(p-k)!}{(p+1)!}.$$

Thus, the flat payoff averages model-switch contributions over both endpoint and intermediate hybrid covariate vectors, whereas $\Lambda$ in Eq. (S3) uses only the two endpoints. Consequently, $\Lambda^{\text{flat}}(\pi)$ and $\Lambda$ need not coincide. Ordinary Shapley efficiency further gives

$$\sum_{j=1}^{p} \Gamma_j^{\text{flat}}(\pi) = \Delta - \Lambda^{\text{flat}}(\pi), \qquad \text{[S7]}$$

so, whenever $\Lambda^{\text{flat}}(\pi) \neq \Lambda$, the flat covariate-switch payoffs do not sum to $\Gamma = \Delta - \Lambda$.

A simple example shows that this discrepancy can occur. We let $p = 2$, take $P_B$ and $P_A$ to be point masses at $(0, 0)$ and $(1, 1)$, respectively, and let $f_B \equiv 0$ and $f_A(x) = x_1 x_2$, for which the coupling is unique. Then $\Delta = 1$ and

$$\Lambda = \Gamma = \tfrac{1}{2}.$$

In the flat three-player game, the model-switch marginal contribution is zero at the hybrid covariate vectors $(0, 0)$, $(1, 0)$, and $(0, 1)$ and equals one at $(1, 1)$. Since the Shapley weight of the predecessor coalition $\{C_1, C_2\}$ is $1/3$,

$$\Lambda^{\text{flat}} = \tfrac{1}{3} \neq \tfrac{1}{2} = \Lambda,$$

and Eq. (S7) gives

$$\Gamma_1^{\text{flat}} + \Gamma_2^{\text{flat}} = \tfrac{2}{3} \neq \tfrac{1}{2} = \Gamma.$$

Thus, a flat Shapley allocation need not preserve the aggregate structure–composition split.

### S3. Existence, Uniqueness, and Admissibility of the MSD Coupling

Because $P_A$ and $P_B$ have finite second moments, we define

$$s_j^2 := \tfrac{1}{2}\{(\Sigma_{X_A})_{jj} + (\Sigma_{X_B})_{jj}\}, \qquad J := \{j \in [p] : s_j > 0\}, \tag{S8}$$

and

$$c(x_A, x_B) := \sum_{j \in J} \left( \frac{x_{Aj} - x_{Bj}}{s_j} \right)^2 . \tag{S9}$$

The MSD coupling is a minimizer of the expected switching cost,

$$\pi^\star \in \arg\min_{\pi} \mathbb{E}_\pi\{c(X_A, X_B)\}, \tag{S10}$$

where the minimum is taken over all couplings $\pi$ of $P_A$ and $P_B$, and multiple minimizers, if any, are resolved by a prespecified deterministic tie-breaking rule. For $j \notin J$, $X_{Aj}$ and $X_{Bj}$ are almost surely constant, so a coupling of $P_A$ and $P_B$ is determined by its restriction to the coordinates in $J$. After standardizing the coordinates in $J$ by $x_j \mapsto x_j/s_j$, Eq. (S10) is equivalent to the Kantorovich optimal-transport problem on $\mathbb{R}^{|J|}$ with squared Euclidean ground cost, and finite second moments ensure the existence of a minimizer (Peyré and Cuturi, 2019; Santambrogio, 2015; Villani, 2009). A standard sufficient condition for uniqueness is absolute continuity, with respect to Lebesgue measure on $\mathbb{R}^{|J|}$, of either standardized endpoint distribution restricted to the coordinates in $J$ (Brenier, 1991; McCann, 1995).

**Proposition 2 (Admissibility).** *On any domain where the MSD minimizer in* Eq. (S10) *is unique, the coupling rule $(P_A, P_B) \mapsto \pi^\star$ is admissible. Under nonuniqueness, admissibility holds if the tie-breaking rule is equivariant to group exchange and common covariate relabeling.*

*Proof.* Suppose first that $P_A = P_B = P$. The identity coupling $\mathcal{L}(X, X)$ has zero cost, so any minimizer has zero cost, which implies $X_{Aj} = X_{Bj}$ almost surely for every $j \in J$. For $j \notin J$, the common marginal coordinate is constant. Hence $X_A = X_B$ almost surely, and the identity coupling is the unique minimizer, so identity at coincidence holds without tie-breaking.

We let $\tau(x_A, x_B) := (x_B, x_A)$. Because Eq. (S8)–Eq. (S9) are invariant to group exchange, $\pi \mapsto \tau_{\#}\pi$ preserves the objective and maps the coupling set for $(P_A, P_B)$ bijectively onto that for $(P_B, P_A)$. Thus, unique minimizers are transposed under group exchange, and the same holds under nonuniqueness when the tie-breaking rule is transposition equivariant. Likewise, a common permutation of the coordinates maps feasible couplings bijectively to the relabeled problem and leaves the standardized cost unchanged, so unique minimizers, or minimizers chosen by a coordinate-equivariant tie-breaking rule, are relabeled in the same way. These are the three admissibility requirements. □

The maximum-entropy tie-breaking rule of the empirical implementation, given in the Computational Procedures of the main text, satisfies this equivariance requirement. Exchanging the groups transposes the cost matrix $(C_{ii'})$ and hence the set $\widehat{\Pi}_{\mathrm{MSD}}$ of empirical minimizers, and the entropy $\mathcal{H}$ is invariant under transposition, so the selected minimizer $\hat{\pi}^\star$ is transposed. A common relabeling of the covariates leaves the cost matrix, and hence $\hat{\pi}^\star$, unchanged.

### S4. Independence and Entropically Regularized Couplings

Under the independence coupling $\pi^{\mathrm{ind}} := P_A \otimes P_B$,

$$Q^{\pi^{\mathrm{ind}}}_{\Omega(H)} = P_{A,H} \otimes P_{B,H^c}, \qquad H \subseteq [p],$$

so the switched and retained blocks are independent, whereas the dependence within each block is preserved. For $\varepsilon > 0$, the entropically regularized coupling is

$$\pi^\varepsilon := \arg\min_{\pi} \Big[\mathbb{E}_\pi\{c(X_A, X_B)\} + \varepsilon \, \mathrm{KL}(\pi \,\|\, P_A \otimes P_B)\Big],$$

where KL denotes the Kullback–Leibler divergence and the minimum is taken over all couplings of $P_A$ and $P_B$. Under the stated finite-second-moment conditions, the entropically regularized problem has a unique minimizer (Csiszár, 1975; Peyré and Cuturi, 2019).

**Lemma S1.** *The independence and entropically regularized couplings satisfy conditions (ii) and (iii) of Definition 1 of the main text but violate condition (i): if $P_A = P_B = P$ and $P$ is not a point mass, neither equals the identity coupling $\pi^{\mathrm{id}} := \mathcal{L}(X, X)$ with $X \sim P$. Hence, neither coupling rule is admissible on a domain containing such a pair $(P, P)$.*

*Proof.* Exchanging $P_A$ and $P_B$ transposes $P_A \otimes P_B$, and a common relabeling of the coordinates relabels it in the same way. Both the switching cost $c$ and $\mathrm{KL}(\cdot \parallel P_A \otimes P_B)$ are invariant under these operations, and $\pi^\varepsilon$ is unique, so $\pi^\varepsilon$ is transposed and relabeled in the same way.

We now let $P_A = P_B = P$ and $\mathcal{I} := \{(x,x) : x \in \mathbb{R}^p\}$, so that $\pi^{\mathrm{id}}(\mathcal{I}) = 1$. If $P$ is not a point mass, then $q := (P \otimes P)(\mathcal{I}^c) > 0$, so $\pi^{\mathrm{ind}} = P \otimes P \neq \pi^{\mathrm{id}}$.

Suppose that $\pi^\varepsilon = \pi^{\mathrm{id}}$. If $\mathrm{KL}(\pi^{\mathrm{id}} \parallel P \otimes P) = \infty$, the objective is infinite at $\pi^{\mathrm{id}}$ but finite at $P \otimes P$, a contradiction. Otherwise, $\pi^{\mathrm{id}} \ll P \otimes P$, so $P = \sum_x p_x \delta_x$ is purely atomic and $\mathrm{KL}(\pi^{\mathrm{id}} \parallel P \otimes P) = -\sum_x p_x \log p_x < \infty$. For $t \in (0,1)$, we let $\pi_t := (1-t)\pi^{\mathrm{id}} + t\, P \otimes P$. Because $c = 0$ on $\mathcal{I}$, $\mathbb{E}_{\pi_t}\{c(X_A, X_B)\} = t\, \mathbb{E}_{P \otimes P}\{c(X_A, X_B)\}$. On $\mathcal{I}^c$, $\pi_t = t\, P \otimes P$, which contributes $qt \log t$ to $\mathrm{KL}(\pi_t \parallel P \otimes P)$, whereas the contribution of $\mathcal{I}$ differs from its value at $t = 0$ by $O(t)$ because $\mathrm{KL}(\pi^{\mathrm{id}} \parallel P \otimes P) < \infty$. Hence, the objective at $\pi_t$ equals its value at $\pi^{\mathrm{id}}$ plus an increment $\varepsilon q t \log t + O(t)$, which is negative for sufficiently small $t$, contradicting the optimality of $\pi^{\mathrm{id}}$. □

The consequences of this violation at coincident covariate distributions are stated in Proposition 3(iv) and proved in Section S5.

### S5. Proof of the Further Properties of MAD

**Proposition 3 (Further properties of MAD).** *MAD satisfies the following properties: (i) coupling invariance of the aggregate effects: $\Lambda$ and $\Gamma$ do not depend on the coupling; (ii) coupling invariance under an additive midpoint predictor: if $\bar{f}(x) = c_0 + \sum_{j=1}^p \psi_j(x_j)$, where $c_0$ is constant and the univariate functions $\psi_1, \ldots, \psi_p$ have finite expectations under $P_A$ and $P_B$, then $\Gamma_j^\pi = \mathbb{E}_{P_A}\{\psi_j(X_j)\} - \mathbb{E}_{P_B}\{\psi_j(X_j)\}$ for any coupling $\pi$ and $j \in [p]$; (iii) null structure effect: if $f_A(x) = f_B(x)$ for $P_A$- and $P_B$-almost every $x$, then $\Lambda = 0$; (iv) null composition effects at coincident distributions: if $P_A = P_B = P$, then $\Gamma = 0$ for any coupling $\pi$; if the coupling is selected by a rule satisfying condition (ii) of Definition 1 of the main text, then $\Gamma_j^\pi = 0$ for every $j \in [p]$; if the rule satisfies condition (i), then $\mathbb{E}_\pi\{\bar{f}(X_{\Omega(H)})\} = \mathbb{E}_P\{\bar{f}(X)\}$ for every $H \subseteq [p]$; and (v) group-exchange equivariance: if the coupling is selected by a rule satisfying condition (ii), exchanging groups $A$ and $B$ reverses the signs of $\Lambda$, $\Gamma$, and each covariate-specific composition effect.*

*Proof.* Part (i) follows immediately from Eq. (S3), which depends only on the endpoint predictor functions and endpoint covariate distributions.

For part (ii), additivity gives, for $H \subseteq [p] \setminus \{j\}$,

$$\mathbb{E}_\pi\{\bar{f}(X_{\Omega(H \cup \{j\})})\} - \mathbb{E}_\pi\{\bar{f}(X_{\Omega(H)})\} = \mathbb{E}_{P_A}\{\psi_j(X_j)\} - \mathbb{E}_{P_B}\{\psi_j(X_j)\},$$

because the two hybrid vectors differ only in coordinate $j$, whose distribution is $P_{A,j}$ under the first and $P_{B,j}$ under the second. The right-hand side does not depend on $H$ or $\pi$, and substitution into Eq. (S4), together with $\sum_{H \subseteq [p] \setminus \{j\}} \kappa_{|H|} = 1$, proves part (ii).

For part (iii), if $f_A(x) = f_B(x)$ for $P_A$- and $P_B$-almost every $x$, then $d = 0$ under both endpoint distributions, and the first identity in Eq. (S3) gives $\Lambda = 0$.

For part (iv), if $P_A = P_B = P$, the second identity in Eq. (S3) gives $\Gamma = 0$ for any coupling $\pi$. Under condition (ii), exchanging the groups leaves $(P, P)$ unchanged, so $\tau_\# \pi = \pi$ for $\tau(x_A, x_B) = (x_B, x_A)$. Hence, with $m^\pi(H) := \mathbb{E}_\pi\{\bar{f}(X_{\Omega(H)})\}$, we have $m^\pi(H) = m^\pi(H^c)$. For $H \subseteq [p] \setminus \{j\}$, we let $H' := ([p] \setminus \{j\}) \setminus H$. Since $(H \cup \{j\})^c = H'$, $H^c = H' \cup \{j\}$, and $\kappa_{|H|} = \kappa_{|H'|}$, reindexing Eq. (S4) gives $\Gamma_j^\pi = -\Gamma_j^\pi$, so $\Gamma_j^\pi = 0$. Under condition (i), $\pi$ is the identity coupling, so $X_{\Omega(H)} = X_A = X_B$ almost surely and $m^\pi(H) = \mathbb{E}_P\{\bar{f}(X)\}$ for every $H \subseteq [p]$.

For part (v), we let tildes denote the quantities after exchanging groups $A$ and $B$. Under condition (ii), transposition equivariance gives $\widetilde{\pi} = \tau_\# \pi$, where $\tau(x_A, x_B) = (x_B, x_A)$. Then $\widetilde{d} = -d$ and $\widetilde{\bar{f}} = \bar{f}$, so Eq. (S3) gives

$$\widetilde{\Lambda} = -\Lambda, \qquad \widetilde{\Gamma} = -\Gamma.$$

Moreover, under the transposed coupling,

$$\widetilde{X}_{\Omega(H)} = X_{\Omega(H^c)}, \qquad H \subseteq [p].$$

For $H \subseteq [p] \setminus \{j\}$, we let $H' := ([p] \setminus \{j\}) \setminus H$, so that $(H \cup \{j\})^c = H'$ and $H^c = H' \cup \{j\}$. Since $\kappa_{|H|} = \kappa_{|H'|}$, reindexing Eq. (S4) gives

$$\widetilde{\Gamma}_j^{\widetilde{\pi}} = -\Gamma_j^\pi, \qquad j \in [p],$$

which proves part (v). □

### S6. Proof of the Reduction to the Blinder–Oaxaca Decomposition

**Proposition 4 (Reduction to the Blinder–Oaxaca decomposition).** *Suppose that $f_g(x) = \alpha_g + x^\top \beta_g$ for $g \in \{A, B\}$, with $\Delta\alpha$, $\Delta\beta_j$, and $\Delta\mu_{X_j}$ as in the Blinder–Oaxaca decomposition of the main text. Then, for any coupling $\pi$,*

$$\Lambda = \Delta\alpha + \sum_{j=1}^p \frac{\mu_{X_{Aj}} + \mu_{X_{Bj}}}{2} \Delta\beta_j, \qquad \Gamma_j^\pi = \Delta\mu_{X_j} \frac{\beta_{Aj} + \beta_{Bj}}{2}, \qquad j \in [p].$$

*Hence, with the componentwise Blinder–Oaxaca terms of the main text evaluated at* $W = \frac{1}{2}I_p$, $\Lambda = \Lambda_0^{\mathrm{BO}} + \sum_{j=1}^{p} \Lambda_j^{\mathrm{BO}}$ *and* $\Gamma_j^{\pi} = \Gamma_j^{\mathrm{BO}}$ *for every* $j \in [p]$.

*Proof.* Since $d(x) = \Delta\alpha + x^{\top}\Delta\beta$, the first identity in Eq. (S3) gives the expression for $\Lambda$. The midpoint predictor

$$\bar{f}(x) = \tfrac{1}{2}(\alpha_A + \alpha_B) + \sum_{j=1}^{p} \tfrac{1}{2}(\beta_{Aj} + \beta_{Bj})x_j$$

is additive, so Proposition 3(ii) with $\psi_j(x_j) = \frac{1}{2}(\beta_{Aj} + \beta_{Bj})x_j$ gives the expression for $\Gamma_j^{\pi}$ for any coupling $\pi$.

At $W = \frac{1}{2}I_p$, equivalently $w_j = \frac{1}{2}$ for every $j \in [p]$, which is the midpoint reference specification of Reimers (1983), the componentwise Blinder–Oaxaca terms (Blinder, 1973; Oaxaca, 1973) are

$$\Lambda_0^{\mathrm{BO}} = \Delta\alpha, \qquad \Lambda_j^{\mathrm{BO}} = \frac{\mu_{X_{Aj}} + \mu_{X_{Bj}}}{2}\Delta\beta_j, \qquad \Gamma_j^{\mathrm{BO}} = \Delta\mu_{X_j}\left(\beta_{Bj} + \frac{1}{2}\Delta\beta_j\right) = \Delta\mu_{X_j}\frac{\beta_{Aj} + \beta_{Bj}}{2},$$

which proves the second statement. □

### S7. Approximate Allocation and Computational Complexity

We use the empirical quantities of the Computational Procedures of the main text. For an empirical coupling $\hat{\pi} \in \mathcal{U}_{n_A,n_B}$, we let

$$\hat{m}_{\hat{\pi}}(H) := \sum_{i=1}^{n_A}\sum_{i'=1}^{n_B} \hat{\pi}_{ii'}\, \hat{\bar{f}}\left(X_{\Omega(H)}(i,i')\right), \qquad H \subseteq [p],$$

denote the internal-game coalition value under $\hat{\pi}$. Conditional on the data and fitted predictor functions, we let $\hat{\Gamma}_j$ and $\hat{\Gamma}_j^{(R)}$ denote the corresponding exact and approximate allocations.

**Approximate allocation.** For a uniformly distributed permutation $\rho$ of $[p]$ and $H \subseteq [p] \setminus \{j\}$, $\Pr\{\mathrm{Pre}_j(\rho) = H\} = \kappa_{|H|}$, the weight of $H$ in the exact allocation. Each summand of the approximate allocation $\hat{\Gamma}_j^{(R)}$ therefore has expectation $\hat{\Gamma}_j$, so $\hat{\Gamma}_j^{(R)}$ is unbiased for the exact allocation under permutation sampling for every fixed $R$ (Castro et al., 2009). Within one permutation, the marginal contributions of the $p$ covariates telescope to $\hat{m}_{\hat{\pi}}([p]) - \hat{m}_{\hat{\pi}}(\emptyset) = \hat{\Gamma}$, which gives $\sum_{j=1}^{p} \hat{\Gamma}_j^{(R)} = \hat{\Gamma}$.

**Adaptive permutation budget.** The budget $R$ may be fixed in advance or selected from pilot permutations. In the adaptive version, we draw $R_0 \geq 2$ independent uniform pilot permutations $\rho_1^{(0)}, \ldots, \rho_{R_0}^{(0)}$, let

$$Z_{mj}^{(0)} := \hat{m}_{\hat{\pi}}\left(\mathrm{Pre}_j(\rho_m^{(0)}) \cup \{j\}\right) - \hat{m}_{\hat{\pi}}\left(\mathrm{Pre}_j(\rho_m^{(0)})\right)$$

denote the marginal contribution of covariate $j$ along pilot permutation $m$, and compute

$$\widetilde{\Gamma}_j := \frac{1}{R_0}\sum_{m=1}^{R_0} Z_{mj}^{(0)}, \qquad \widetilde{\sigma}_j^2 := \frac{1}{R_0 - 1}\sum_{m=1}^{R_0}\left(Z_{mj}^{(0)} - \widetilde{\Gamma}_j\right)^2, \qquad j \in [p].$$

We measure the Monte Carlo error of the vector of approximate allocations by its relative standard error, $\{\sum_j \mathrm{Var}(\hat{\Gamma}_j^{(R)})\}^{1/2}/(\sum_j \hat{\Gamma}_j^2)^{1/2}$, which for a budget $R$ is estimated from the pilot by $\{\sum_j \widetilde{\sigma}_j^2/(R\sum_j \widetilde{\Gamma}_j^2)\}^{1/2}$. Given a target tolerance $\zeta > 0$ and bounds $R_{\min} \leq R_{\max}$, we set

$$R_{\mathrm{req}} := \left\lceil \frac{\sum_{j=1}^{p} \widetilde{\sigma}_j^2}{\zeta^2 \sum_{j=1}^{p} \widetilde{\Gamma}_j^2} \right\rceil, \qquad R_{\mathrm{ad}} := \min\{R_{\max}, \max(R_{\min}, R_{\mathrm{req}})\},$$

where $R_{\mathrm{req}}$ is the smallest budget at which the estimated relative standard error does not exceed $\zeta$. If $\sum_{j=1}^{p} \widetilde{\Gamma}_j^2$ is numerically zero, the relative criterion is undefined, and we set $R_{\mathrm{ad}} := R_{\min}$ when $\sum_{j=1}^{p} \widetilde{\sigma}_j^2$ is also numerically zero and $R_{\mathrm{ad}} := R_{\max}$ otherwise.

The allocation is then computed from $R_{\mathrm{ad}}$ new permutations sampled independently of the pilot, and the pilot contributions are not reused. Conditional on the pilot, $R_{\mathrm{ad}}$ is fixed and the new permutations are independent and uniformly distributed, so the unbiasedness and adding-up properties above hold as for a fixed budget. The criterion targets an estimated relative Monte Carlo precision of the vector of covariate-specific composition effects.

**Computational complexity.** We let $L_{\hat{\pi}} := \#\{(i,i') : \hat{\pi}_{ii'} > 0\}$ denote the support size of $\hat{\pi}$ and $C_{\mathrm{pred}}$ the cost of evaluating $\hat{\bar{f}}$ at one $p$-dimensional covariate vector. For coupling construction, the empirical MSD and entropically regularized couplings require the switching-cost matrix $(C_{ii'})$, whose construction takes $O(n_A n_B p)$ operations and $O(n_A n_B)$ storage. When $n_A = n_B = n$, the empirical MSD problem is a linear assignment problem, solvable in $O(n^3)$ operations, and a unique optimal assignment gives $L_{\hat{\pi}^\star} = n$. The empirical independence coupling requires no optimization and can be represented implicitly in

$O(1)$ time and storage. For the empirical entropically regularized coupling, $T$ Sinkhorn iterations require $O(Tn_An_B)$ operations, and the solution has the form $\operatorname{diag}(a)\,(e^{-C_{ii'}/\varepsilon})\,\operatorname{diag}(b)$ with positive vectors $a$ and $b$ (Peyré and Cuturi, 2019). Hence $L_{\hat{\pi}^{\text{ind}}} = L_{\hat{\pi}^{\varepsilon}} = n_An_B$.

For allocation, each coalition value $\hat{m}_{\hat{\pi}}(H)$ requires $L_{\hat{\pi}}$ evaluations of $\hat{\bar{f}}$. The exact allocation computes and stores all $2^p$ coalition values, requiring $O(2^pL_{\hat{\pi}}C_{\text{pred}})$ operations for their evaluation, $O(p2^p)$ operations for the Shapley weighting, and $O(2^p)$ storage. With a fixed budget $R$, the approximate allocation evaluates $p+1$ coalition values along each sampled permutation, requiring $O(RpL_{\hat{\pi}}C_{\text{pred}})$ operations and $O(p)$ storage for the running averages; with an adaptive budget, $R$ is replaced by $R_0 + R_{\text{ad}}$. Substituting $L_{\hat{\pi}^\star} = n$ and $L_{\hat{\pi}^{\text{ind}}} = L_{\hat{\pi}^{\varepsilon}} = n^2$ gives the allocation costs in the complexity table of the main text.

### S8. Training Specifications of the Predictive Models

For each group $g \in \{A, B\}$, predictive-model training used a sample of size $n_{\text{train}} = 1000$ drawn independently of the sample used for MAD. Tuning was performed within the training sample; unspecified settings used software defaults.

**Linear regression.** Linear regression was fitted by least squares with an intercept and all $p$ covariates as main effects, without nonlinear transformations or interaction terms.

**Gradient boosting machines.** Gradient boosting was implemented with `XGBoost` (Chen and Guestrin, 2016) and `LightGBM` (Ke et al., 2017) using squared-error loss. Maximum depth $\{1, 2, 3, 4\}$ and learning rate $\{0.01, 0.05\}$ were selected by 10-fold cross-validation; boosting iterations were selected by early stopping with a maximum of 2,000 rounds and patience 50. The selected configuration was refitted on all $n_{\text{train}} = 1000$ observations.

**Deep neural networks.** Deep neural networks were implemented with `Keras` (Chollet et al., 2015) and `Torch` (Paszke et al., 2019) using the same training procedure. Both used two ReLU hidden layers, a linear output activation, and mean squared error loss. Hidden-layer widths were selected from $\{(p, \lceil p/2 \rceil), (2p, p), (4p, 2p)\}$ and dropout from $\{0, 0.15, 0.3\}$ using two held-out folds of a five-fold partition of the training sample. For each of these two folds, predictors were standardized using the remaining four folds as the fitting subset, and each network was trained with Adam (Kingma and Ba, 2015) (learning rate 0.01, batch size 100) for at most 500 epochs with early stopping on the held-out fold (patience 50) and evaluated at its best validation epoch. The configuration with the smallest mean validation RMSE across the two held-out folds was selected, and the final predictor averaged the predictions of its two best-epoch networks, so that every training observation contributed to the final model without an additional fit.

### S9. Population Reference Quantities for the Decomposition Study

Except for the LA aggregate references given below, the population references are replicate-specific because the coefficient and mean-shift directions and the correlation matrices are regenerated in each decomposition-study replicate. Replicate superscripts are suppressed below. We let $f_g^0 = \eta_g^{(\ell)}$ and $\bar{f}^0 = (f_A^0 + f_B^0)/2$.

**Population MSD coupling.** For the decomposition study,

$$X_A \sim N_p(\delta, \Sigma_{X_A}), \qquad X_B \sim N_p(0, \Sigma_{X_B}),$$

where $\Sigma_{X_A}$ and $\Sigma_{X_B}$ are the unstructured correlation matrices specified in the main text. Because both have unit diagonal, $s_j = 1$ in Eq. (S8). The population MSD coupling is the Gaussian quadratic optimal-transport coupling (Gelbrich, 1990; Peyré and Cuturi, 2019),

$$X_A = \delta + A_{\text{OT}}X_B, \qquad A_{\text{OT}} = \Sigma_{X_B}^{-1/2}\bigl(\Sigma_{X_B}^{1/2}\Sigma_{X_A}\Sigma_{X_B}^{1/2}\bigr)^{1/2}\Sigma_{X_B}^{-1/2}.$$

Hence every hybrid vector $X_{\Omega(H)}$ is Gaussian.

**Aggregate reference quantities.** With $\mu_{r|s}^0 := \mathbb{E}_{P_s}\{f_r^0(X)\}$ for $r, s \in \{A, B\}$,

$$\Delta^0 = \mu_{A|A}^0 - \mu_{B|B}^0, \qquad \Lambda^0 = \tfrac{1}{2}\bigl(\mu_{A|A}^0 + \mu_{A|B}^0 - \mu_{B|A}^0 - \mu_{B|B}^0\bigr),$$

$$\Gamma^0 = \tfrac{1}{2}\bigl(\mu_{A|A}^0 + \mu_{B|A}^0 - \mu_{A|B}^0 - \mu_{B|B}^0\bigr), \qquad \Delta^0 = \Lambda^0 + \Gamma^0.$$

We write $\mu_{X_A} = \delta$ and $\mu_{X_B} = 0$, let $K = 2p/5$, and define

$$q_{g|s,j} := \mathbb{E}_{P_s}\{\phi_{r_g(j)}(X_j)\}, \qquad m_{g|s,k} := \mathbb{E}_{P_s}\{\chi_{g,k}(X)\},$$

for $g, s \in \{A, B\}$, $j \in [p]$, and $k = 1, \dots, K$, and let $m_{g|s} = (m_{g|s,1}, \dots, m_{g|s,K})^\top$. Then

$$\mu^{0,\mathrm{LA}}_{A|s} = \alpha + (\beta+\gamma)^\top \mu_{X_s}, \qquad \mu^{0,\mathrm{LA}}_{B|s} = \beta^\top \mu_{X_s},$$

$$\mu^{0,\mathrm{LM}}_{A|s} = \alpha + (\beta+\gamma)^\top \mu_{X_s} + (\theta+\xi)^\top m_{A|s}, \qquad \mu^{0,\mathrm{LM}}_{B|s} = \beta^\top \mu_{X_s} + \theta^\top m_{B|s},$$

$$\mu^{0,\mathrm{NA}}_{A|s} = \alpha + \sum_{j=1}^p (\beta_j+\gamma_j) q_{A|s,j}, \qquad \mu^{0,\mathrm{NA}}_{B|s} = \sum_{j=1}^p \beta_j q_{B|s,j},$$

$$\mu^{0,\mathrm{NM}}_{A|s} = \alpha + \sum_{j=1}^p (\beta_j+\gamma_j) q_{A|s,j} + (\theta+\xi)^\top m_{A|s}, \qquad \mu^{0,\mathrm{NM}}_{B|s} = \sum_{j=1}^p \beta_j q_{B|s,j} + \theta^\top m_{B|s}.$$

Because every marginal under $P_B$ is $N(0,1)$ and each $\phi_k$ is centered to have mean zero under $N(0,1)$, $q_{A|B,j} = q_{B|B,j} = 0$ for all $j$. The interaction expectations are obtained from Gaussian second- and third-order moments (Isserlis, 1918). In particular, if $X \sim N_p(\mu, \Sigma)$,

$$\mathbb{E}(X_a X_b) = \mu_a \mu_b + \Sigma_{ab},$$

and

$$\mathbb{E}(X_a X_b X_c) = \mu_a \mu_b \mu_c + \mu_a \Sigma_{bc} + \mu_b \Sigma_{ac} + \mu_c \Sigma_{ab}.$$

The nonlinear expectations for $\phi_1$, $\phi_2$, and $\phi_4$ are evaluated in closed form, whereas $\phi_3$ is evaluated by one-dimensional numerical integration.

For LA,

$$\beta^\top \delta = \gamma^\top \delta = \frac{p}{4},$$

so

$$(\mu^0_{A|A}, \mu^0_{A|B}, \mu^0_{B|A}, \mu^0_{B|B}) = \left(1 + \frac{p}{2}, 1, \frac{p}{4}, 0\right),$$

and

$$(\Delta^0, \Lambda^0, \Gamma^0) = \left(1 + \frac{p}{2}, 1 + \frac{p}{8}, \frac{3p}{8}\right).$$

For LM, NA, and NM, the references are evaluated replicate by replicate from the formulas above.

**Covariate-specific reference quantities.** The reference $\Gamma^0_j$ is $\Gamma^{\pi^\star}_j$ in Eq. (S4) evaluated with $\bar{f}^0$, where $\pi^\star$ is the population MSD coupling above. For LA and NA, $\bar{f}^0(x) = \alpha/2 + \sum_{j=1}^p \psi_j(x_j)$ is additive, with $\psi_j(t) = (\beta_j + \gamma_j/2)\, t$ for LA and $\psi_j(t) = \frac{1}{2}\{(\beta_j+\gamma_j)\phi_{r_A(j)}(t) + \beta_j \phi_{r_B(j)}(t)\}$ for NA, so Proposition 3(ii) gives, for LA,

$$\Gamma^0_j = \left(\beta_j + \frac{\gamma_j}{2}\right)\delta_j = \frac{p}{4}\left(\nu_{1j} + \frac{1}{2}\nu_{2j}\right)(\nu_{1j} + \nu_{2j} + \nu_{3j}),$$

with

$$\sum_{j=1}^p \Gamma^0_j = \beta^\top\delta + \frac{1}{2}\gamma^\top\delta = \frac{3p}{8} = \Gamma^0,$$

and, for NA,

$$\Gamma^0_j = \frac{\beta_j+\gamma_j}{2}\left[\mathbb{E}_{P_A}\{\phi_{r_A(j)}(X_j)\} - \mathbb{E}_{P_B}\{\phi_{r_A(j)}(X_j)\}\right] + \frac{\beta_j}{2}\left[\mathbb{E}_{P_A}\{\phi_{r_B(j)}(X_j)\} - \mathbb{E}_{P_B}\{\phi_{r_B(j)}(X_j)\}\right].$$

For LM and NM, the midpoint predictors are

$$\bar{f}^0_{\mathrm{LM}}(x) = \frac{\alpha}{2} + \sum_{j=1}^p \left(\beta_j + \frac{\gamma_j}{2}\right) x_j + \frac{1}{2}\sum_{k=1}^K \left[(\theta_k+\xi_k)\chi_{A,k}(x) + \theta_k \chi_{B,k}(x)\right],$$

and

$$\bar{f}^0_{\mathrm{NM}}(x) = \frac{\alpha}{2} + \frac{1}{2}\sum_{j=1}^p \left[(\beta_j+\gamma_j)\phi_{r_A(j)}(x_j) + \beta_j \phi_{r_B(j)}(x_j)\right] + \frac{1}{2}\sum_{k=1}^K \left[(\theta_k+\xi_k)\chi_{A,k}(x) + \theta_k \chi_{B,k}(x)\right].$$

Substitution into Eq. (S4) reduces the interaction contributions to second- and third-order moments of the Gaussian hybrid vectors. These moments are evaluated analytically under the population MSD coupling for each replicate.

### S10. Coalition Deviation under Approximate Allocation

In the coupling study, the coalition deviation of an empirical coupling $\hat{\pi}$ is

$$D_{\hat{\pi}} := \left\{ 2^{-p} \sum_{H \subseteq [p]} \left[ \hat{m}_{\hat{\pi}}(H) - \hat{m}_{\hat{\pi}}(\emptyset) \right]^2 \right\}^{1/2},$$

with $\hat{m}_{\hat{\pi}}(H)$ as in Section S7. For $p \in \{5, 10, 15\}$, where the exact allocation is used, $D_{\hat{\pi}}$ is computed from all $2^p$ coalition values. For $p = 20$, where the approximate allocation is used, it is estimated from the same sampled permutations $\rho_1, \ldots, \rho_R$. With $H_{mk}$ denoting the set of the first $k$ covariates in $\rho_m$, for $k = 0, \ldots, p$, we use

$$\widehat{D}_{\hat{\pi},R} := \left[ \sum_{k=0}^{p} \frac{\binom{p}{k}}{2^p} \frac{1}{R} \sum_{m=1}^{R} \{ \hat{m}_{\hat{\pi}}(H_{mk}) - \hat{m}_{\hat{\pi}}(\emptyset) \}^2 \right]^{1/2}.$$

For each $k$, $H_{mk}$ is uniform over subsets of size $k$, so $\widehat{D}^2_{\hat{\pi},R}$ is unbiased for $D^2_{\hat{\pi}}$. The calculation reuses the allocation paths and requires no additional coalition evaluations.

### S11. Details of the Real Data Application

**Data processing.** We used the closed-reference operational taxonomic unit table of the AGP (McDonald et al., 2018), in which reads trimmed to 100 nucleotides were clustered at 97% similarity against the Greengenes 13_8 reference (DeSantis et al., 2006; McDonald et al., 2012), without rarefaction. Bacterial counts were aggregated to the genus level after bracketed genus names were merged with their unbracketed counterparts and reads without a genus assignment were removed. We retained fecal samples from US and UK residents aged 20–70 years with a recorded sex, a BMI of at least 15 and below 60, and at least 10,000 reads, and kept the sample with the largest number of reads for each participant. Genera with a mean relative abundance of at least 0.001 among the resulting 6,673 participants were retained, and the analysis was then restricted to males aged 30–70 years. Zero counts were replaced by 0.5 before the CLR transformation.

**Training procedures.** Each predictive model was fitted separately to each country sample, with BMI as the response and the $p = 53$ CLR coordinates as covariates; unspecified settings used software defaults. Linear regression was fitted by least squares with an intercept and all CLR coordinates as main effects; because the CLR coordinates sum to zero, its coefficients are determined only up to a common additive constant, which leaves the fitted predictor on the CLR hyperplane unchanged. Gradient boosting with `XGBoost` (Chen and Guestrin, 2016) and `LightGBM` (Ke et al., 2017) used the squared-error loss; maximum depth $\{1, 2, 3, 4\}$ and learning rate $\eta \in \{0.01, 0.05\}$ were selected by two repeats of 10-fold cross-validation, boosting iterations were selected by early stopping with a maximum of 2,000 rounds and patience 50 at $\eta = 0.05$ and 10,000 rounds and patience 250 at $\eta = 0.01$, and the selected configuration was refitted on all observations. Deep neural networks with `Keras` (Chollet et al., 2015) and `Torch` (Paszke et al., 2019) used two ReLU hidden layers, a linear output activation, and the mean squared error loss. Hidden-layer widths $\{(\lceil p/2 \rceil, \lceil p/4 \rceil), (p, \lceil p/2 \rceil), (2p, p)\}$ and dropout $\{0, 0.1, 0.2, 0.3\}$ were selected by five-fold cross-validation. Within each fold, predictors and BMI were standardized using the fitting subset, and each network was trained with Adam (Kingma and Ba, 2015) (learning rate 0.01, batch size 100) for at most 500 epochs with early stopping on the held-out fold (patience 50) and retained at its best validation epoch; its predictions were transformed back to the BMI scale and shifted to match the mean BMI of the fitting subset. The final predictor averaged the five networks of the selected configuration and was shifted to match the mean BMI of the country sample.

**Predictive performance.** Predictive performance was assessed by the out-of-fold root mean squared error (RMSE) in 10-fold cross-validation within each country, with all tuning carried out within the training folds, and was averaged over the two countries. The average RMSEs (kg/m$^2$) of linear regression, `XGBoost`, `LightGBM`, `Keras`, and `Torch` were 4.179, 4.189, 4.161, 4.180, and 4.162, respectively, and `LightGBM`, with the smallest average RMSE, was selected. Applying its tuning procedure to each full country sample selected trees of depth one with $\eta = 0.05$ and 365 (UK) and 314 (US) boosting iterations.

**Evaluation at hybrid covariate vectors.** We let $\Pi_{\mathrm{CLR}} := I_p - p^{-1}\mathbf{1}_p\mathbf{1}_p^\top$ denote the orthogonal projection onto the CLR hyperplane $\{x \in \mathbb{R}^p : \mathbf{1}_p^\top x = 0\}$, and we applied MAD to $\tilde{f}_g(x) := \hat{f}_g(\Pi_{\mathrm{CLR}} x)$ for $g \in \{A, B\}$. Because $\Pi_{\mathrm{CLR}} x = x$ for every observed CLR vector, $\tilde{f}_g$ and $\hat{f}_g$ coincide on both samples, so that $\Delta$, $\Lambda$, and $\Gamma$ are unchanged and only the internal-game coalition values at intermediate hybrid covariate vectors are affected.

**Coupling and allocation.** The empirical MSD coupling was supported on $n_A + n_B - 1 = 2{,}593$ pairs. The adaptive budget of Section S7 used $R_0 = 500$, $R_{\min} = 500$, $R_{\max} = 100{,}000$, and $\zeta = 0.0045$, slightly below the target of 0.005, which gave $R_{\mathrm{ad}} = 646$ and an achieved relative Monte Carlo standard error of 0.0045. Table S1 lists the genus-specific composition effects of all 53 genera; their Monte Carlo standard errors were at most $1.1 \times 10^{-4}$.

**Table S1. Genus-specific composition effects $\Gamma_j$ (kg/m$^2$, UK minus US) of the 53 genera, in decreasing order of $|\Gamma_j|$ down the left and then the right column; $\sum_{j=1}^{53} \Gamma_j = \Gamma = -0.100$.**

| Genus | $\Gamma_j$ | Genus | $\Gamma_j$ |
|---|---|---|---|
| *Corynebacterium* | 0.0765 | *Bacillus* | 0.0028 |
| *Finegoldia* | −0.0648 | *Megasphaera* | −0.0021 |
| *Bifidobacterium* | 0.0439 | *Turicibacter* | 0.0021 |
| *Clostridium* | −0.0345 | *Stenotrophomonas* | −0.0020 |
| *Haemophilus* | −0.0281 | *Eubacterium* | −0.0018 |
| *Akkermansia* | −0.0214 | *Desulfovibrio* | 0.0016 |
| *Butyricimonas* | 0.0202 | *Anaerococcus* | −0.0016 |
| *Dorea* | −0.0154 | *Enterobacter* | 0.0015 |
| *Collinsella* | −0.0151 | *Serratia* | 0.0015 |
| *Catenibacterium* | 0.0109 | *Lachnospira* | −0.0012 |
| *Ruminococcus* | −0.0096 | *Paraprevotella* | 0.0012 |
| *Prevotella* | −0.0082 | *Streptococcus* | −0.0011 |
| *Blautia* | −0.0082 | *Morganella* | −0.0010 |
| *Lachnobacterium* | −0.0078 | *Bilophila* | −0.0006 |
| *Peptoniphilus* | −0.0063 | *Staphylococcus* | −0.0006 |
| *Porphyromonas* | −0.0063 | *Fusobacterium* | −0.0006 |
| *Dialister* | −0.0061 | *Sutterella* | 0.0005 |
| *Roseburia* | −0.0060 | *Coprococcus* | −0.0004 |
| *Anaerostipes* | −0.0055 | *Pseudomonas* | −0.0003 |
| *Odoribacter* | 0.0045 | *Bacteroides* | 0.0003 |
| *Lactobacillus* | −0.0042 | *Veillonella* | −0.0003 |
| *Listeria* | −0.0041 | WAL_1855D | 0.0002 |
| *Succinivibrio* | 0.0036 | *Acinetobacter* | 0.0002 |
| *Parabacteroides* | −0.0035 | *Brevundimonas* | −0.0001 |
| *Faecalibacterium* | 0.0033 | *Comamonas* | 0.0001 |
| *Oscillospira* | −0.0032 | *Proteus* | −0.0001 |
| *Phascolarctobacterium* | −0.0030 | | |
